# Rapid Caspian sea level decline requires dynamic spatio-temporal conservation planning to keep biodiversity protection measures relevant

**DOI:** 10.1101/2024.11.08.622693

**Authors:** Rebecca Court, Matteo Lattuada, Nataliya Shumeyko, Mirgaliy Baimukanov, Tariel Eybatov, Altynay Kaidarova, Elchin V. Mamedov, Eldar Rustamov, Aselle Tasmagambetova, Matthias Prange, Thomas Wilke, Christopher Hassall, Simon J. Goodman

**Author notes:** **Correspondence:** Rebecca Court, Simon J. Goodman, Email:, School of Biology, Faculty of Biological Sciences, University of Leeds, Leeds, LS2 9JT, UK. School of Earth and Environment, Faculty of Environment, University of Leeds, Leeds, LS2 9JT, UK. Plant Ecology, Institute of Ecology, Technische Universität Berlin, 12165 Berlin, Germany (Matteo Lattuada).

## Abstract

The Caspian Sea is the world’s largest landlocked waterbody, providing habitat for hundreds of endemic and migratory species, along with ecosystem services that sustain millions of people. Global warming is projected to drive declines in water levels of up to 21 m by 2100. Using geospatial analyses, we assessed the impact of sea level decline on habitats, protected areas, and human infrastructure. We show that a water level decline of just 5–10 m will critically disrupt key ecosystems, reduce existing marine protected area coverage by up to 94%, and render billions of dollars of civil and industrial infrastructure obsolete. Urgent action is needed to replace traditional static conservation planning with a pre-emptive, dynamic approach that allows protected areas to track shifting ecosystems. This will be essential to help the Caspian Sea’s endemic biodiversity adapt to these changes, and avoid conflicts with mitigation efforts directed at protecting human activity.

**Russian abstract:** **Абстракт**

Каспийское море – крупнейший в мире внутренний водоем, обеспечивающий среду обитания для сотен эндемичных и мигрирующих видов, а также экосистемные услуги, обеспечивающие жизнедеятельность миллионов людей. Согласно прогнозам, глобальное потепление приведет к снижению уровня воды до 21 м к 2100 году. С помощью геопространственного анализа проведена оценка влияния снижения уровня моря на места обитания, охраняемые территории и объекты инфраструктуры. Мы показываем, что снижение уровня воды всего на 5–10 м приведет к критическим нарушениям в ключевых экосистемах, сократит охват существующих морских охраняемых районов до 94% и приведет к выводу из эксплуатации гражданской и промышленной инфраструктуры стоимостью в миллиарды долларов. Необходимы безотлагательные меры по замене традиционного статичного планирования природоохранной деятельности упреждающим, динамичным подходом, который позволит корректировать границы охраняемых районов вслед за смещением экосистем. Это необходимо для адаптации эндемичного биоразнообразия Каспийского моря к изменениям, а также для того, чтобы усилия по его сохранению не вступали в конфликт с мерами по защите деятельности человека.

## Introduction

Climate change is driving environmental crises for landlocked water bodies and rivers globally, with profound consequences for biodiversity, agriculture, fisheries, infrastructure and other economic activity, as well as human health and wellbeing (Woolway et al., 2020). Increased rates of evaporation and changes to precipitation patterns are leading to declines in the levels of lakes and landlocked seas (Dai et al., 2018; W. Wang et al., 2018; Yao et al., 2023), meaning that current spatial conservation measures need to be adapted to remain relevant as key habitats shift to track changing water levels (Markovic et al., 2014; Parks et al., 2022; Pittock et al., 2008). Significant changes in lake/inland sea ice phenologies are also expected (Li et al., 2022). Adapting protected areas for these ecosystems to make them fit for purpose in future will need to account for conflicts with concomitant actions to mitigate impacts on civil and industrial infrastructure and economic activity. However, the systematic evaluations needed to balance conservation and socioeconomic interests in mitigation planning are missing for most major inland waterbodies. Here we assess vulnerability to sea level decline for key ecosystems, protected areas and human infrastructure in the Caspian Sea, the world’s largest landlocked waterbody, and its relevance to developing a pre-emptive, temporally dynamic, spatial marine conservation strategy for the region.

Today, the Caspian Sea (Figure 1) extends approximately 1150km by 450km, and is bordered by five countries: Azerbaijan, Iran, Kazakhstan, Russia, and Turkmenistan. Covering an area of about 387,000 km^2^, it has three sub-basins: the northern Caspian has average water depths of 5m, while the middle and southern basins reach 788m, and more than 1000m respectively. The Caspian Sea receives water from around 130 rivers, with more than 80% of the inflow via the Volga and Ural rivers in the north (Leroy et al., 2020; Rodionov, 1994).

**Figure 1.**
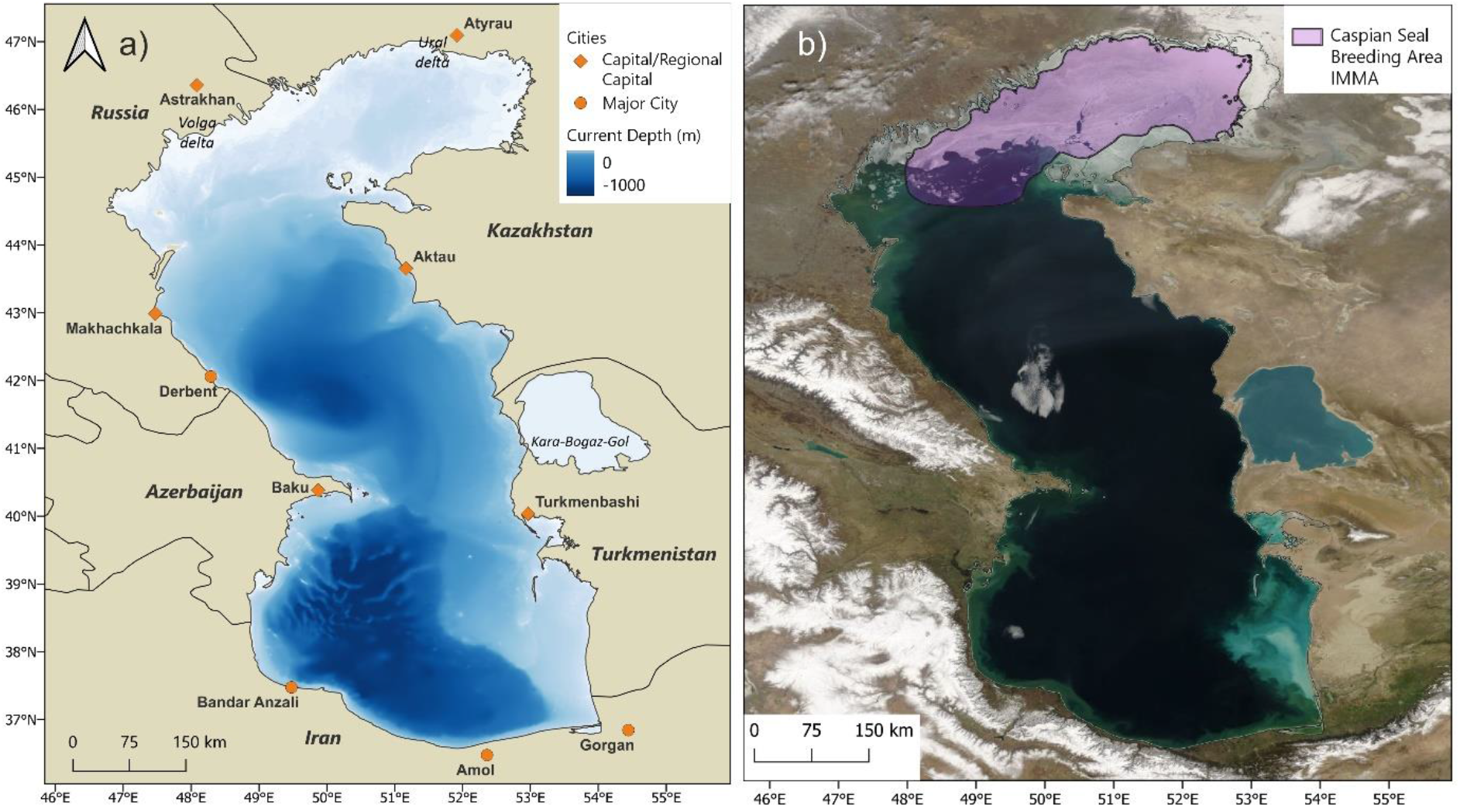
Left, bathymetry of the Caspian Sea with a datum of -27.5m against global mean sea level. Right, NASA MODIS satellite image of the Caspian Sea region showing typical winter ice coverage of the northern Caspian basin, which is the breeding habitat for the endemic Caspian seal (Pusa caspica). The polygon indicates the Caspian seal breeding Important Marine Mammal Area (IMMA; see methods).

The depth and surface area of the Caspian Sea have varied over time, with regressions of up to 90m and transgressions of 50m during the Quaternary (Krijgsman et al., 2019). At the beginning of the 20th century, Caspian sea level (CSL) was approximately -25.8m below global mean sea level. It then decreased by 2m, stabilised during the 1950s and 1960s, before reducing rapidly between 1970 and 1977, reaching its lowest level in 100 years at -29m. The level then rose to -26.5m by 1995, subsequently declining again by around 2m (Chen et al., 2017; Lahijani, Leroy, et al., 2023). The 20^th^ Century variations are attributed to changes in water inflow rates (due to natural factors and human abstraction) and precipitation, and their balance against evaporation. The current declining trend is linked to global warming driving reduced precipitation, and a sharp increase in sea surface temperature (SST) and evaporation (Arpe et al., 2014; Chen et al., 2017; J. Wang et al., 2018). CSL declined by −6.07 ± 0.26 cm per year between 2002 and 2015 (Chen et al., 2017), a negative rate 20 times greater than global sea level increase. Declines as high as 30cm per year have been measured since 2020 (CASPCOM, 2024). Early modelling studies predicted climate-driven CSL declines of around 5m by 2100 (Elguindi & Giorgi, 2006; Renssen et al., 2007), but more recent projections predict reductions of 9 to 21m by the end of the century under medium to high emission scenarios (Koriche, Singarayer, et al., 2021; Nandini-Weiss et al., 2019; Samant & Prange, 2023).

The Caspian Sea is of high importance for biodiversity in Central Asia and Europe, supporting more than 300 species of endemic invertebrate, and 76 endemic fish (Karpinsky, 2010), as well as the endemic Caspian seal (*Pusa caspica*) (Goodman & Dmitrieva, 2016). The coastal margins provide habitats for globally significant populations of water and migratory birds, with major flyways connecting Europe, Asia, and Southern Africa (Veen et al., 2005). The Caspian Sea has experienced substantial anthropogenic impacts and stressors including habitat destruction, pollution, introduction of invasive species and overexploitation of natural resources (Lattuada et al., 2019; Mammadov et al., 2016); leading to extinctions and displacement of native fauna (Lattuada et al., 2020), and collapse of commercial fisheries (Strukova et al., 2016) and the seal population (Härkönen et al., 2012). CSL decline threatens already impacted ecosystems, the winter sea ice breeding habitat of Caspian seals in the northern basin (Wilson et al., 2017), and the stability of regional climate (Koriche, Nandini-Weiss, et al., 2021). While concerns over the consequences of CSL decline for biodiversity have been raised previously (Prange et al., 2020; Samant & Prange, 2023), little systematic evaluation has been carried out on vulnerabilities of ecosystems and conservation protections arising from Caspian sea level decline.

More than 15 million people live around the Caspian coast (United Nations Environment Programme, 2011). The projected sea level decline presents major economic and societal implications. Shipping routes, civil and industrial infrastructure, including ports and hydrocarbon production facilities, could become obsolete or unviable, while industrial and artisanal fisheries face severe disruption. As with the Aral Sea, increased exposure to pollutant contaminated sediment dust, arising from desiccation, could cause serious human health effects (Bennion et al., 2007). Adverse societal consequences from changes in CSL have previously been reported (Rodionov, 1994) but are largely unrecognised in current policy and conservation spatial planning (Prange et al., 2020).

Quantitative assessments of vulnerabilities are required for Caspian countries to develop effective mitigation plans that can help sustain ecosystem services and functions from impacts arising from CSL declines. To address this gap, we first quantify the extent of coastal recession in the northeastern Caspian Sea from 2001 to 2022. Next, we estimate how CSL decline will reduce coverage of key ecosystems and existing spatial conservation designations in the Caspian region under different future decline scenarios, and evaluate vulnerabilities of human infrastructure and marine areas important for economic activity. We then suggest policy priorities for effective, integrated, and sustainable responses to the environmental pressures that will soon unfold in the region.

## Methods

### Data sources

Spatial data were compiled from open access sources. All spatial data processing and analyses were conducted using QGIS 3.22.12 (QGIS, 2023).

A Caspian Sea boundary polygon was constructed using data from the Global Administrative Areas 2015 (v2.8) dataset (http://purl.stanford.edu/zb452vm0926). Gridded bathymetric data was obtained from the General Bathymetric Chart of the Oceans repository at a spatial resolution of 15 arc seconds (equivalent to less than 1km at 40ºN; GEBCO; www.gebco.net/data_and_products/gridded_bathymetry_data/), and clipped to the Caspian Sea boundary, representing the 2010 coastline at datum of -27.5m below mean global sea level (Leroy *et al*., 2020).

Data on key ecosystems, ecologically important areas and current spatial conservation designations within the Caspian region were compiled as follows: i) Ecologically or Biologically Significant Marine Areas (EBSAs; n=15; www.cbd.int/ebsa/); ii) Important Marine Mammal Areas (IMMAs; n=3; www.marinemammalhabitat.org/immas/); iii) Caspian Ecoregions (n=10; a synthesis of distinct Caspian Sea habitat types based on ecological and physical parameters as defined by (Fendereski et al., 2014) (See supplementary Table 1)); iv) Seasonal ranges of Caspian sturgeon species (Karpinsky, 2010); v) Marine Protected Areas (n=16) and Terrestrial and Inland Waters Protected Areas, (n=12), listed in the World Database on Protected Areas (WDPA; www.protectedplanet.net/en/thematic-areas/wdpa), and records from local nature protection agencies. Additional curation of names, designation, and polygon boundaries for protected areas is described in Supplementary Table 2.

EBSAs, designated under the Convention on Biological Diversity, and IMMAs, designated by the IUCN Joint SSC-WCPA Marine Mammal Protected Areas Task Force, are not legally defined protected areas. Rather they are areas of high biodiversity importance identified by expert working groups and are intended to inform spatial planning decisions by policy makers. They may encompass areas recognised as ecologically significant under other designations such as Ramsar sites or national classifications. Areas listed in WDPA may have a formal status under a host country’s national legislation, but with varying levels of protection and enforcement.

Locations of key Caspian seal haul-out sites were taken from the Caspian Sea IMMA documents. These sites had been previously categorised into 1) Currently used sites; 2) Established seal habitat that is no longer used regularly or by a significant number of seals; and 3) A known historical area of seal habitat that is no longer used at all.

Caspian Sea oil and gas infrastructure (on and offshore production and processing facilities, pipelines and ports) and settlements were georeferenced from map data at the University of Texas at Austin GeoData service (https://geodata.lib.utexas.edu/catalog/princeton-736666054), and supplemented with literature and Google Earth searches.

Monthly vessel density data (vessel hours per km^2^ per month) within the Caspian Sea for the calendar year of 2022 was obtained from Global Maritime Traffic (https://globalmaritimetraffic.org/gmtds.html). ‘No data’ values were set to zero, and monthly values summed to give annual values, before being log10 transformed.

### Estimation of shoreline recession in the northeastern Caspian Sea (2001-2022)

The northeastern Caspian (Figure 2) is an area with extremely shallow bathymetry (typically less than 1m), consisting of extensive reed beds, sandbanks, muddy shoals, and water channels. It provides important habitats for water and migratory birds, fish spawning areas, as well as moulting and resting sites for Caspian seals. To document ecosystem transition due to sea level change, satellite images (250m per pixel) were sourced from NASA Worldview (2023, www.worldview.earthdata.nasa.gov/), for March and October of each year between 2001 and 2022. The first cloud-free image closest to the 15^th^ day of each month was selected for analysis. In QGIS, for each image, within a polygon with bounds 47.12° N, 53.43° E to 45.27° N, 52.05° E, the coastline was digitised manually to distinguish separate land and water polygons. The area (km^2^) for each part was then calculated using the QGIS polygon area tool.

**Figure 2.**
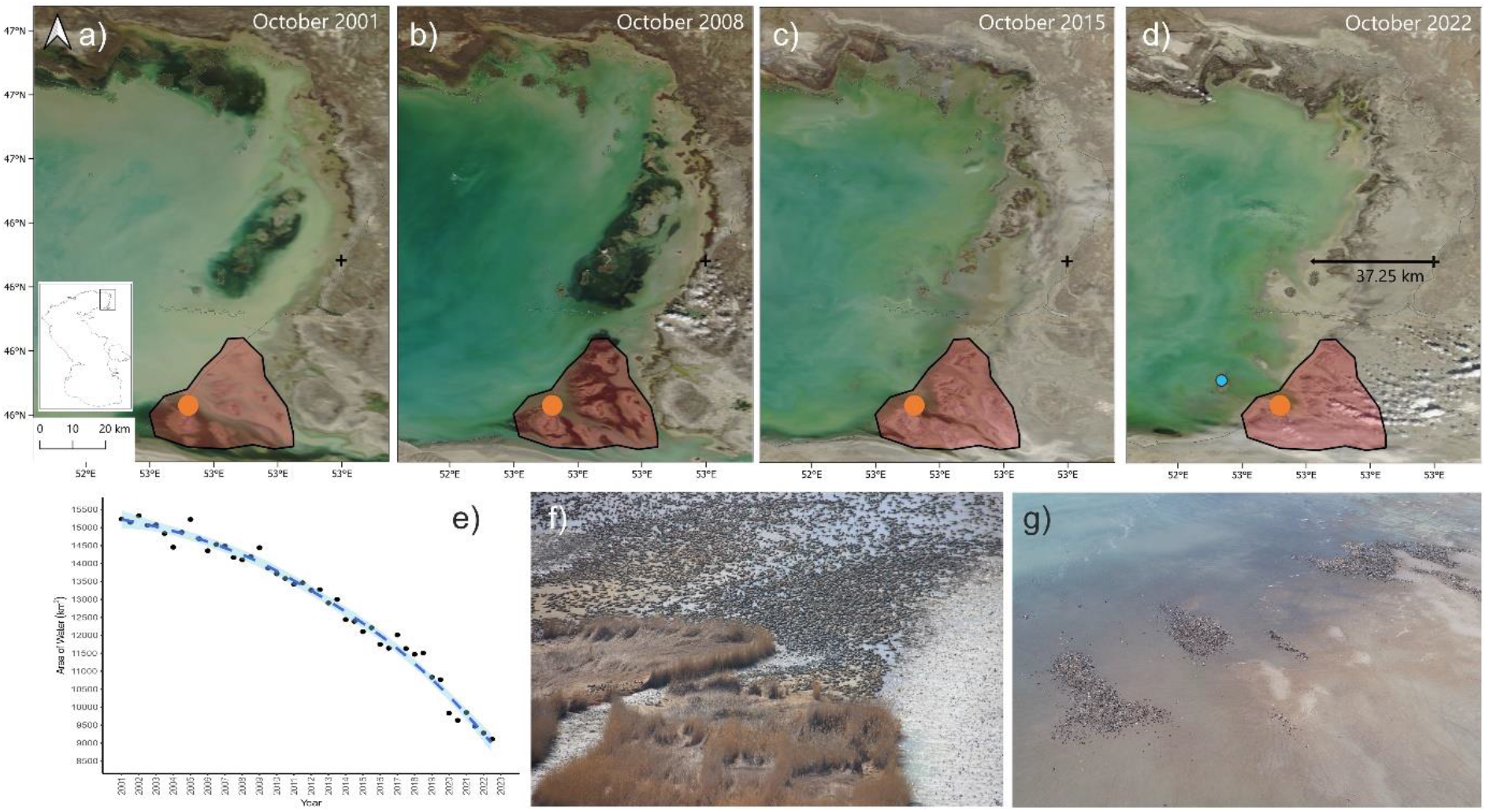
Coastal regression in the northeastern Caspian Sea. a to d) Time series of MODIS satellite images from 2001 to 2022 showing sea level recession; the polygon indicates boundary of Komsomol Bay EBSA. The orange point marks the location of Caspian seal moulting aggregations in April 2011, and the blue point indicates the location of moulting aggregations in April 2022. e) Time series of water area within the focal polygon (area shown in MODIS image) from 2001-2022. f) Dense Caspian seal moulting aggregations in Komsomol Bay, April 2011 (Photo: Lilia Dmitrieva, University of Leeds). g) Moulting aggregations of Caspian seals on newly emergent islands April 2022 (Photo: Assel Baimukanova, Institute of Hydrobiology and Ecology).

### Calculating decline scenario bathymetry

Nandini-Weiss et al. (2019) project sea level decline for the Caspian of up to 18m by 2100. We defined eight sea level decline scenarios in increments of 2.5m (−2.5m, -5m, -7.5m, -10m,

-12.5m, -15m, -17.5m, and -18m decline) relative to the -27.5m datum. Individual scenarios were generated by reclassifying the GEBCO bathymetry raster as 0 or 1 for cells above/below the specified depth of a decline scenario (e.g. -30m for a 2.5m decline). The reclassified raster was multiplied by the original bathymetry values, and the new sea level (e.g., -30m) was added to reach the true depth (where 0m is the surface).

### Reduction in area of marine habitat and conservation zones

For each individual marine zone, the proportional reduction in area covered by water was calculated under the eight sea level decline scenarios, relative to coverage at the -27.5m datum. First the bathymetry scenario rasters were converted to polygons and clipped to the Caspian Sea boundary polygon. Proportional coverage of conservation areas under each decline scenario was then measured using the ‘Overlap Analysis’ tool which calculates area and percent cover of overlaying layers. This returned the area of each conservation site under each decline scenario which were then divided by the original site area.

### Changes in distance to shore

The nearest distance to the shoreline (km) for human settlement and infrastructure georeferenced points was calculated under each decline scenario using the ‘v.distance’ tool in QGIS GRASS. For terrestrial coastal protected areas, distance to shore was calculated from the centroid of the area polygon.

## Results

### Sea level recession in the northeastern Caspian Sea 2001-2022

The focal polygon for sea level recession in the northeastern Caspian Sea covered 21901 km^2^, with 15194 km^2^ of water in 2001 (Figure 2). By October 2022, the area covered by water decreased by 39% to 9197 km^2^, at approximately 300 km^2^ per year (Figure 2e). Relative to the MODIS datum coastline, the shore receded by at least 37.25 km between 2001 and 2022. By 2022, most of the area within the Komsomol Bay EBSA boundary, recognised as an important spring moulting site for Caspian seals (Figure 2f), had become desiccated, and seals could no longer access islands for moulting. Aerial surveys and field observations during spring 2018-2023 indicate moulting seals have shifted to newly emergent islands to the north, but with lower densities than observed in Komsomol Bay (Baimukanov et al 2022; Figure 2g).

### Overview of sea level decline scenarios across Caspian Sea region

A 2.5m decline leads to a significant loss of water coverage around the margins of the northern basin (Figure 3, Supplementary Figure 1). Following a 5m decline, approximately 77,000 km^2^ of current water area (around 20% of the present Caspian Sea surface) will become land, with the most affected areas being the northern basin, Kara-Bogaz-Gol, and Turkmenistan shoreline (Figure 3, Supplementary Figure 1). Under the most extreme decline scenario of 18m (Figure 3; Supplementary Figures 1a to 1h), as much as 143,000 km^2^ of water could be desertified, a loss of 37% relative to the current Caspian Sea area.

**Figure 3.**
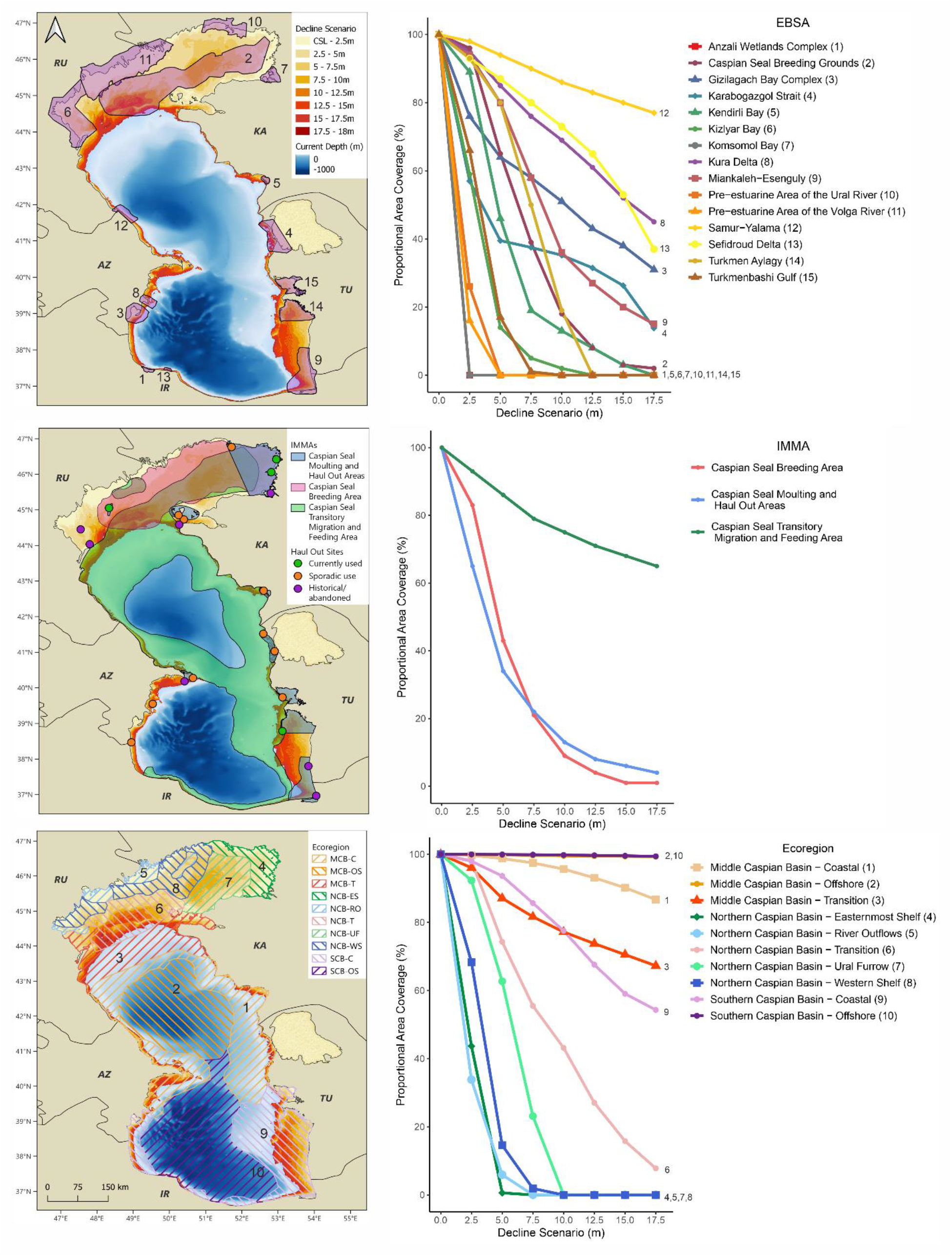
Area reductions arising from different sea level decline scenarios on currently designated Ecological, or Biological Significant Areas (EBSAs; top); Important Marine Mammal Areas (IMMAs; middle); and Caspian Ecoregions (bottom). Left column: maps of decline scenario overlap with spatial area designations. Right column: charts showing proportional reduction in sea area covered by current spatial area designations under the different decline scenarios.

### Reduction of Caspian ecoregions and areas with ecological significance

Most ecoregions and all area designations with recognised ecological significance (EBSAs, IMMAs) would experience increasing reductions in marine coverage (i.e., conversion to land) as sea level declines (Figure 3; Supplementary Tables 3, 4, 5). Under a decline of just 5m, 7 of 15 EBSAs would experience proportional coverage reductions exceeding 50%, with 4 completely desiccated, and similar outcomes for 2 of 3 IMMAs. Among ecoregions, 3 of 5 in the northern Caspian would experience reductions exceeding 80% under a 5m decline. With a 10m decline 5 EBSAs and 4 ecoregions would be completely eliminated, while IMMAs would see reductions ranging from 25% to more than 80% (Figure 3; Supplementary Tables 3, 4, 5).

### Caspian seal breeding and sturgeon habitats

Sea level declines of 2.5m to 5m will make all current and historical seal haul-out sites inaccessible to seals (Figure 3). Shifts in haul-out site use due to sea level recession can already be observed in the northeastern Caspian Sea (Figure 2). The current Caspian seal breeding area in the northern Caspian is extremely vulnerable to sea level reduction. The area of the breeding IMMA covered by water and winter sea ice, and therefore accessible to seals, could be reduced by around 57% with a decline of just 5m, but this could be as much as 81%, if the deeper sub-basin of the northern Caspian becomes isolated from the main portion of the Caspian Sea by a land barrier (Figure 3, 4; Supplementary Table 4).

**Figure 4.**
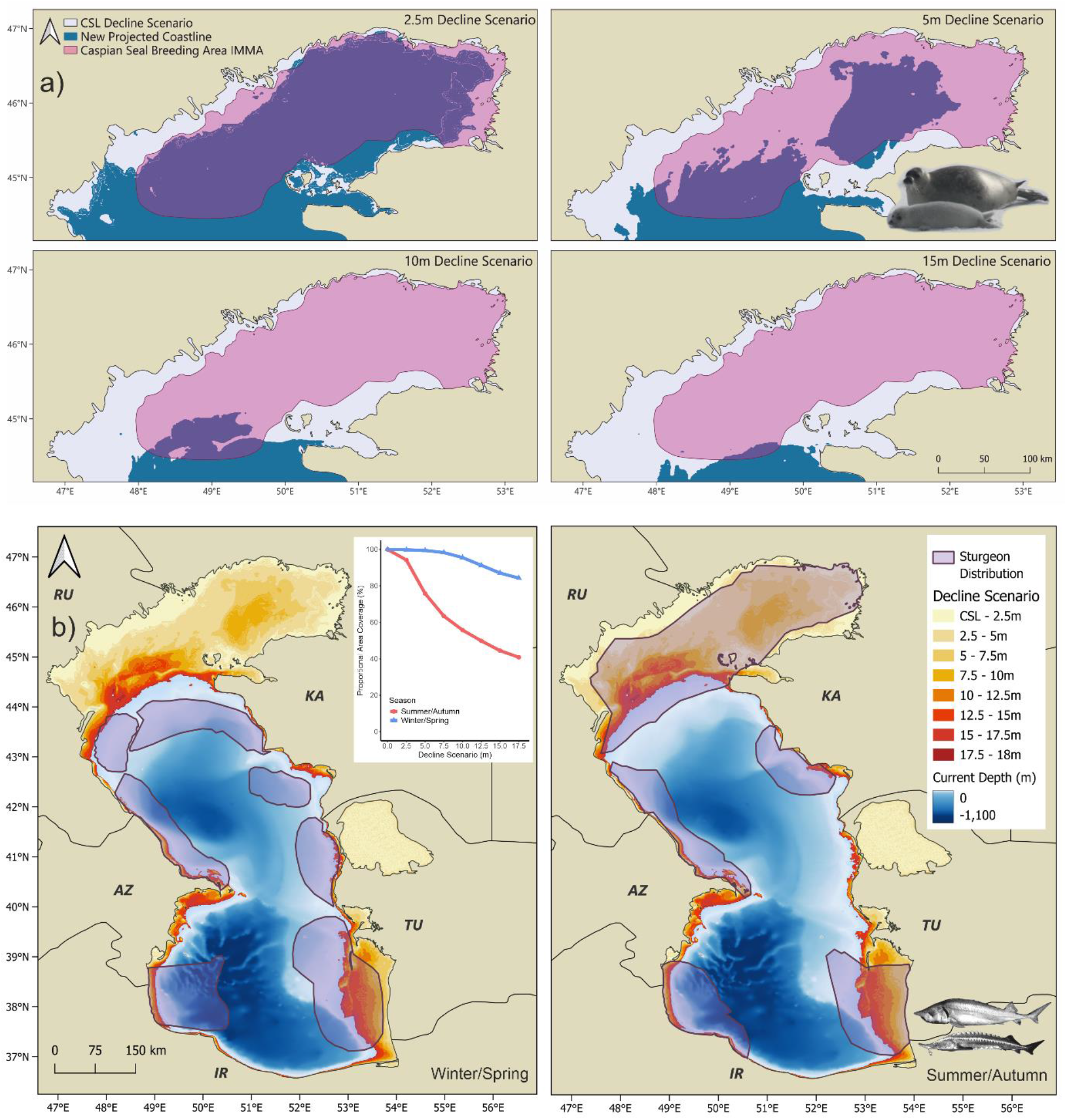
Maps showing: a) Sea area reduction from the Caspian seal breeding IMMA, under 2.5m, 5m, 10m and 15m decline scenarios; and b) Summer/Autumn and Winter/Spring sturgeon ranges under estimated CSL decline scenarios; inset graph quantifies percentage area reduction for each season group.

The summer/autumn ranges of Caspian sturgeon species overlap considerably with shallow water areas, and could experience a 25% reduction with a 5m decline and around a 45% reduction with a 10m decline (Figure 4; Supplementary Table 6). Sturgeon winter/spring ranges are less dependent on shallow water areas, reducing by around 5% at 10m CLS decline.

### Effects on current marine and coastal terrestrial protected areas

All but one WDPA-listed marine area would see coverage reductions exceeding 50% at 5m, with 11 of 16 protected areas transitioning completely to land. Declines of 10m and above eliminate marine coverage for most currently designated conservation areas (Figures 5a,b; Supplementary Table 7). The current total coverage of Caspian Sea marine protected areas is 16.8% of the sea area. A 5m decline would decreases coverage to 7%, and a 10m decline to just 1% (Figure 5c).

**Figure 5.**
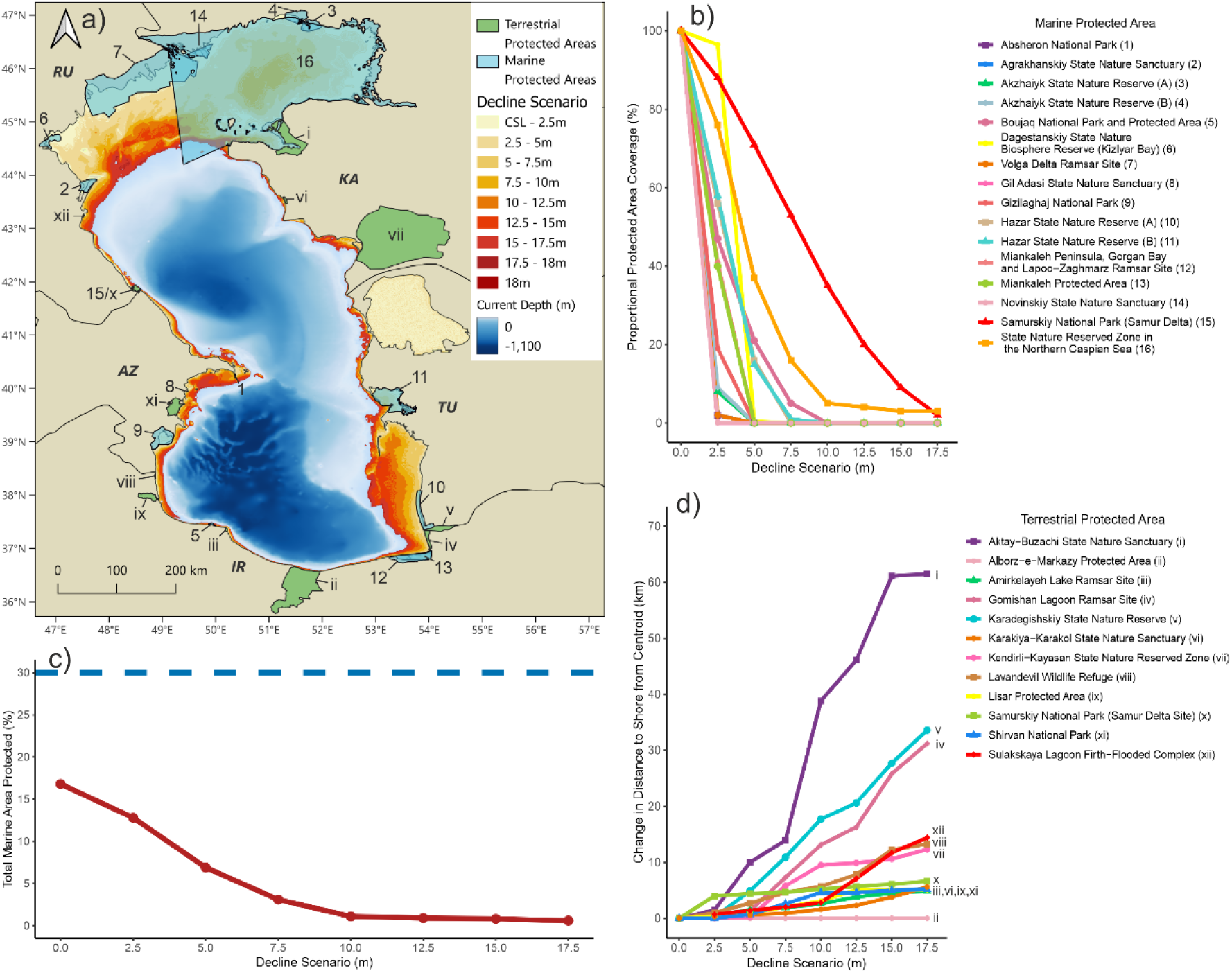
Area reductions arising from different sea level decline scenarios on current areas with legislative environmental protections listed in the World Database on Protected Areas (WDPA); a) Map of decline scenario overlap with spatial area designations; b) Plot showing proportional reduction in sea area covered by current spatial area designations under the different decline scenarios; c) Decline in overall marine protected area coverage relative to Caspian Sea area for the -27.5m datum, dashed reference line indicates the Kunming-Montreal Global Biodiversity Framework (GBF) target of 30% area coverage of inland water, coastal, and marine areas; d) Increase in distance to shore for Caspian coastal terrestrial protected areas, under sea level decline scenarios of 2.5m to 17.5m relative to the study datum.

Under a 5m decline, coastal terrestrial protected areas (shore-based ecosystems that depend on the sea for their function) will experience a multi-kilometre increase in their distance to shore (Figure 5d; Supplementary Table 8). Under a 10m decline, 3 of 12 sites will experience an increase of more than 10 km, increasing to 6 sites with a 15m decline. The greatest increase in distance to shore would be expected for the Aktay-Buzachi State Nature Sanctuary in Kazakhstan, with an increase of 61.7km under the 18m decline scenario.

### Human infrastructure and activity

Coastal settlements in all Caspian countries will experience increases in distance to shore, requiring major adaptation of civil infrastructure for declines of 2.5m and above (Figure 6; Supplementary Table 9). Coastal settlements in the northern Caspian basin, within Kazakhstan and Russia, will experience the greatest increase in their distance to the Caspian shoreline (Figure 6b). Under a 10m decline scenario, the mean increase in distance to shore for Russian and Kazakh settlements is 44 km (range: 18.8-102.2 km) and 89 km (range: 0.87-

**Figure 6.**
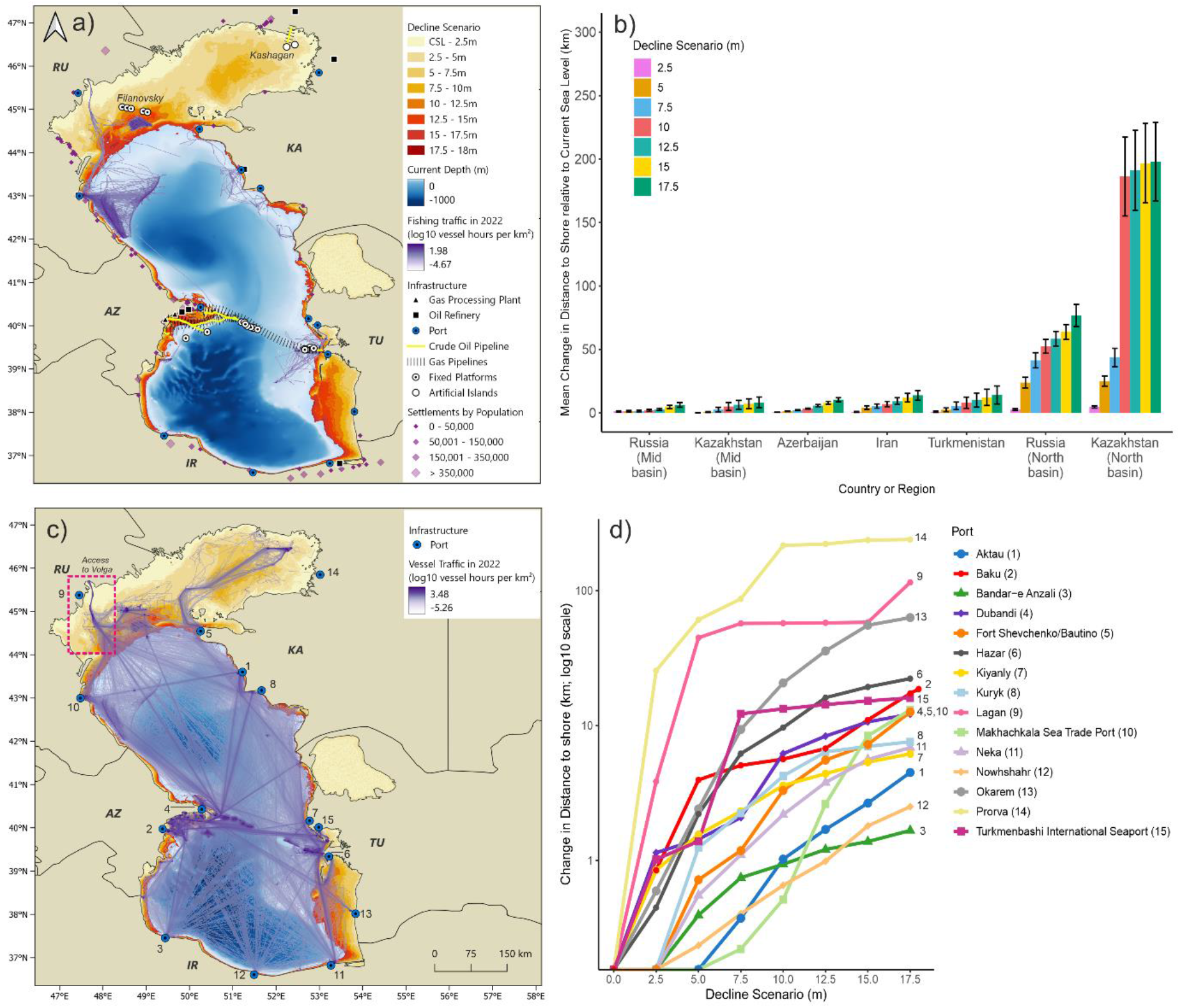
Impact of decline scenarios on human civil and industrial infrastructure and activity. a) Map showing locations of coastal settlements, industrial infrastructure and commercial fishing activity relative to decline scenarios; b) Bar chart summarising mean increase in distance to coast for settlements relative to the -27.5m datum (error bars show standard deviation; values for individual settlements can be found in Supplementary Tables 9, 10), in different Caspian coastal regions; c) Map showing shipping density and port locations relative to decline scenarios; d) Plot showing increase in distance to coast relative to the -27.5m datum for ports.

259.7 km), respectively.

More than one third (8 out of 22) of all industrial infrastructure facilities included in this study, comprising ports, oil refineries and gas processing plants, will experience an increase greater than 5 km in their distance to the Caspian shoreline, under a 10m sea level decline scenario (Figure 6; Supplementary Table 10). Ports that are critical for trade or industrial logistics will be impacted in all countries, with Baku (Azerbaijan), Anzali (Iran), and Aktau (Kazakhstan), experiencing increases in distance to shore of 1 km or more, while Turkmenbashi (Turkmenistan), and Lagan (Russia; planned future site), could see increases of 16 km and 115 km respectively (Figure 6d).

The Kashagan (Kazakhstan) and Filanovsky (Russia) oil fields in the northern Caspian are two of the region’s most important hydrocarbon production sites. Production currently occurs at offshore installations, with ship-based logistics (Figure 6a, b), but these will become landlocked under 5 to 15m CSL declines. Access to the Caspian Sea for international shipping via the Volga River will also be critically impacted by a decline of 5m (Figure 6c), with major consequences for navigation and trade.

Industrial fishing for kilka (*Clupeonella* spp.) is presently focused on relatively deep waters off the coast of Dagestan, but one fishing ground on the boundary between the northern and mid Caspian basin could disappear with declines above 5m (Figure 6a). Artisanal fisheries, which primarily use shallower waters, will also be heavily impacted, particularly in the northern Caspian Sea, where declines of 5m can be expected to eradicate most fishing activity.

## Discussion

We made the first quantitative assessment of impacts from projected 21^st^ Century sea level decline on Caspian Sea ecosystems and spatial conservation designations. Our analyses suggest that CSL declines of 5 to 10m, as projected for medium emissions scenarios by the end of this century, will cause losses exceeding 80% of current extent for many ecosystem types and ecologically important areas. These declines will also eliminate or substantially reduce the marine coverage of most current spatial conservation designations in the Caspian Sea. In parallel, 5 to 10m declines will have dramatic consequences for coastal communities, industrial infrastructure and economic activity, as settlements, ports, and offshore energy installations become stranded tens or hundreds of kilometres from new shorelines.

The sensitivity of Caspian region protected areas to climate change has previously been recognised (Lahijani, Azizpour, et al., 2023; Tulloch et al., 2020; Zheng et al., 2021), but there has been no previous quantitative evaluation of their vulnerability to CSL decline. We find that without mitigation, marine protected area coverage will reduce from 16.8% of the Caspian Sea extent at the -27.5m datum to 7% and 1% for 5m and 10m decline scenarios respectively. The high sensitivity of current spatial conservation designations arises because they are aligned with ecologically significant areas, and regions of high habitat diversity around coasts and transition zones into deeper waters in the northern Caspian basin, and other shallow shelf areas. The northern basin has a mean depth of just 5m, and although it accounts for only 1% of the total Caspian Sea water volume, with an extent of approximately 115,000 km^2^, it represents one-third of the total sea surface area.

As CSL decline proceeds, current marine areas will shift from permanent water coverage, to transitory inundation, and eventual desiccation, with water associated vegetation such as reed beds and *Salicornia* (a genus of succulent, halophytic, flowering plants) transitioning to desert or steppe (Lahijani, Azizpour, et al., 2023). Shallow shelf and transition zone ecosystems will be greatly reduced, without replacement, since areas with remaining water coverage have steeper bathymetry. Habitats for endemic benthos will be displaced and diminished as water levels reduce because they are constrained to specific bathymetry ranges, substrate types, water oxygen levels, and experience competition from invasive species (Lattuada et al., 2020). New terrestrial habitats such as emergent islands may develop, but the succession process and their suitability to sustain displaced species is poorly studied. Current coastal terrestrial protected areas will experience multi-kilometre increases in their distance to shore, meaning the ecosystems they encompass (lagoons, reed beds, salt marshes and beaches) will also transition to steppe and desert. Overall, there will be significant loss of internationally important habitats (recognised by their EBSA and Ramsar designations) for fish, birds and benthos, with potential for cascading effects throughout Caspian Sea ecosystems and beyond, for example due to loss of fish spawning areas and habitats supporting migratory birds.

Such changes are already visible around the margins of the northern Caspian Sea and other areas with shallow bathymetry such as Turkmenbashi Bay (Turkmenistan), Gorgan Bay and Gomishan Lagoon (Iran), and the Ghizil Agaj Bay (Azerbaijan) (Lahijani, Azizpour, et al., 2023). In our focal northeastern Caspian study zone water coverage has decreased by approximately 6,000km^2^ (39%), between 2001 and 2022, leading to displacement of Caspian seal moulting and haul-out sites in the Komsomol Bay EBSA, which is now nearly completely desiccated. Recession is also visible within the Volga and Ural deltas, including the 3410km^2^ Akzhayik State Nature Reserve close to Atyrau. Caspian seal haul-out sites on the Zyud-Vestovye Shalygas Islands, located in the buffer zone of the Akzhaiyk State Nature Reserve, have also been lost (Baimukanov et al., 2022).

The endangered Caspian seal (Goodman & Dmitrieva, 2016), is vulnerable to loss of its present breeding area. The species pups on the winter ice sheet that currently forms in the northern basin December to March each year (Wilson et al., 2017). A 5m decline would lead to reductions between 57% and 81% of the current breeding IMMA. How climate warming and sea level reduction will interact to influence the Caspian ice regime, and to what extent winter sea ice might persist around the northern coast of a reduced Caspian Sea is poorly understood (Naurozbayeva et al., 2023; Tamura-Wicks et al., 2015), as is whether Caspian seals can successfully adapt to terrestrial breeding in the absence of ice, for example on newly emergent islands. The northern Caspian basin also provides important foraging areas for Caspian seals (Dmitrieva et al., 2016), and other culturally and economically important species such as Caspian sturgeon (Pueppke et al., 2023). Loss of the area may compromise access to important sturgeon spawning rivers such as the Ural, and spawning habitat of the commercially important common kilka (*Clupeonella caspia*).

CSL decline will also cause significant impacts for human wellbeing, infrastructure, and economic activity. The economic cost of disruption could exceed tens of billions of dollars per year. Millions of tons of cargo are shipped through the Volga River each year, the only external maritime connection, to and from Caspian (Akbulaev & Bayramli, 2020). Access to the Volga could be compromised by declines of as little as 5m, with dredging already implemented to keep passage for larger vessels open (Akundov, 2024). Similarly, operators of offshore oil production facilities in the northern Caspian basin are dredging channels to maintain shipping logistics (North Caspian Operating Company, 2022), and operations at Aktau port and the intake for the city’s water desalination plant have been disrupted due to declining water levels (Amangeldina, 2024).

For artisanal and coastal fishing communities in the northern basin, economies are likely to collapse entirely as fishing grounds disappear. Alternative incomes sources from aquaculture, horticulture, and ecotourism (Svolkinas et al., 2023), may also become inviable posing risks to the social integrity of communities. Commercial fisheries will shift to deeper waters where possible, with consequent changes in maritime zones of jurisdiction and exclusive fishing rights. However, they may be vulnerable to economic losses if fish stocks fall due to loss of spawning habitats and other CSL decline related impacts.

Ecosystem services provided by the Caspian Sea to the surrounding region will also be altered by sea level reductions. The cumulative effects of decreased ice cover, increased surface water temperatures, and changes in sea level will cause alterations in mixing-regimes and deep ventilation, with consequences for nutrient cycling (Woolway et al., 2020). Physical atmospheric changes in precipitation and moisture may increase aridity of the regional climate, exacerbating water scarcity across central Asia (Koriche, Nandini-Weiss, et al., 2021). Desiccation of inland lakes can also pose health risks to local inhabitants. Previous evaporation events following restriction of the connection from the Caspian into Kara-Bogaz-Gol, and in the Aral Sea, led to toxic dust and salt exposure from the emerged seabed, causing serious concerns for respiratory health and land degradation (Bennion et al., 2007). For similar reasons, along with anticipated harsh climatic conditions, it is unlikely that newly exposed land would be suitable for agriculture (Issanova et al., 2022).

Our assessments are based on projections of CSL decline derived from the latest generation of climate models that suggest reductions in water levels of up to 21m by 2100 under high emission scenarios. Lesser declines (5 to 12m), drying most of the northern Caspian basin will still occur even if global temperature rises are kept below 2ºC under the Paris Climate Accord (Koriche, Singarayer, et al., 2021; Samant & Prange, 2023). Timelines for reductions are uncertain, but levels fell by 1.4m in the 16 years 2006 to 2021, and approximately 75cm 2022 to 2024 (CASPCOM, 2024), a rate that if maintained, imply declines of 17m or more by 2100. Changes in land-use and increases in water abstraction (e.g. for desalination) may compound climate-driven declines (Hutson & Taganova, 2023; Samant & Prange, 2023). The behaviour of potential isolated sub-basin remnants in the northern Caspian Sea is also highly uncertain as this will depend on whether they maintain river connections (e.g. with the Ural), and flow volumes, with implications for ecosystem persistence and conservation management in the early stages of the drying of the northern basin.

We employed bathymetry data with a resolution of 15 arc seconds (equivalent to less than 1km). This is sufficient to assess risks to broadscale ecosystem coverage and large protected areas, but finer scale bathymetry data is required for detailed assessments for specific local areas and individual pieces of infrastructure. We used remaining water coverage as a proxy measure for an area’s ability to sustain habitats and ecosystem processes. However, habitat suitability and ecosystem sustainability may scale non-linearly as water levels decrease, and ecosystem functionality could fail at smaller proportional reductions if environmental factors exceed species tolerances, for example, minimum water depth, water temperature ranges, oxygen and/or nutrient availability, substrate type, or loss of co-dependent species.

### Conclusions and recommendations

Sea level decline will amplify existing anthropogenic threats to Caspian biodiversity (Lattuada et al., 2019). For example, species range shifts may increase species exposure to degraded habitats, existing industrial activities, or other forms of human-wildlife conflict such as fisheries. Similarly, mitigation of CSL decline in relation to human infrastructure and activities has the potential to create new impacts, or conflicts with conservation priorities. Current Caspian Sea spatial conservation designations have varying levels of protections and enforcement, and there are few marine protected areas that prohibit all human economic activity. They will require substantial adaptation to remain relevant and fit for purpose.

Addressing CSL decline impacts on biodiversity will require adaptive, temporally dynamic approaches to spatial conservation planning. Rather than having fixed spatial boundaries, protected areas will need to track shifting species ranges and ecosystems over time, similar to proposals suggested in the context of coral reef conservation (Beger et al., 2022; Buenafe et al., 2023). Planning processes need to be forward-looking, so that areas predicted to host important habitats and ecosystems in future receive pre-emptive protection that can be considered when planning for concomitant mitigation for impacts on human infrastructure and activity. For example, conflicts may arise between ecosystem conservation and relocation/adaptation of ports or shipping routes. Such an approach will be essential to allow temporal and spatial contiguity of ecosystems, and for maintaining commitments under the Kunming-Montreal Global Biodiversity Framework (GBF) such as conserving 30% of inland water, coastal, and marine areas through area-based conservation measures (Secretariat of the Convention on Biological Diversity, 2023).

Legislation such as the Kazakh EcoCode (Parliament of the Republic of Kazakhstan, 2021), in principle allows for some dynamic protections, but Caspian countries may need to revise legal frameworks on protected areas to formalise the status of adaptive as opposed to fixed boundaries. It will also be necessary to collect more comprehensive species and community occurrence data and establish monitoring, to support development of detailed species distribution models for Caspian taxa, evaluations of species vulnerabilities, and predictions of species range shifts through time. Such models will be necessary to inform spatial planning and decisions on other types of conservation interventions such as assisted translocations or ecosystem restoration, coupled with detailed risk assessments for civil and industrial infrastructure, and their associated mitigation plans (Tulloch et al., 2020). Investment to enhance regional expertise in biodiversity and spatial planning should also be prioritised.

Commitment to economic diversification and sustainable development goals will be essential to the survival of coastal communities. Sources of income such as fishing are likely to become unviable for communities in the north. Ensuring human prosperity will be central to maintaining both social stability and sustainable biodiversity (Svolkinas et al., 2023). Communication and consultation with communities will be essential for finding equitable solutions.

Effective responses to CSL declines will require rapid and coordinated transboundary cooperation on governance. Since the end of the Soviet Union in the early 1990s, the Caspian states have made progress on establishing legal frameworks for environmental governance under the Tehran Convention. However, they have often prioritised national economic and political interests over transboundary cooperation to address ecological degradation (Bayramov, 2020). The Caspian Sea serves as an example of the urgent need to address climate change impacts on major landlocked waterbodies worldwide, where pre-emptive, adaptive planning will be required to keep spatial conservation relevant and manage inevitable conflicts with parallel mitigation to protect human populations. Successfully adapting to these impacts in the Caspian region will require a level of transboundary political cooperation beyond that achieved to date. The anticipated scale of environmental disruption in the Caspian Sea will potentially have global consequences, and therefore warrants attention and support from the wider international community.

## Supporting information

Court_et_al_2024_Sup_info_part_1

Court_et_al_2024_Sup_info_part_2

## Acknowledgements

No specific funding was used in the development of this work. Field research by MB in Kazakhstan was supported by the Ministry of Agriculture of the Republic of Kazakhstan (Grant No. BR23591095). We are grateful to the Astrakhanskiy State Nature Biosphere Reserve and Dagestanskiy State Nature Biosphere Reserve for their comments and GIS resources that help support curation of information relating to their protected areas. We thank Caitlin Hayman, Dr Lilia Dmitrieva and Dr Harrison Tan for assistance with georeferencing of Caspian region infrastructure, and Prof Maria Beger for comments on an early draft of the manuscript. We also thank Assel Baimukanova for providing a Russian translation of the abstract, and for comments on the manuscript.

## Ethics statement

No ethical review was required for this work.

## CRediT author contributions

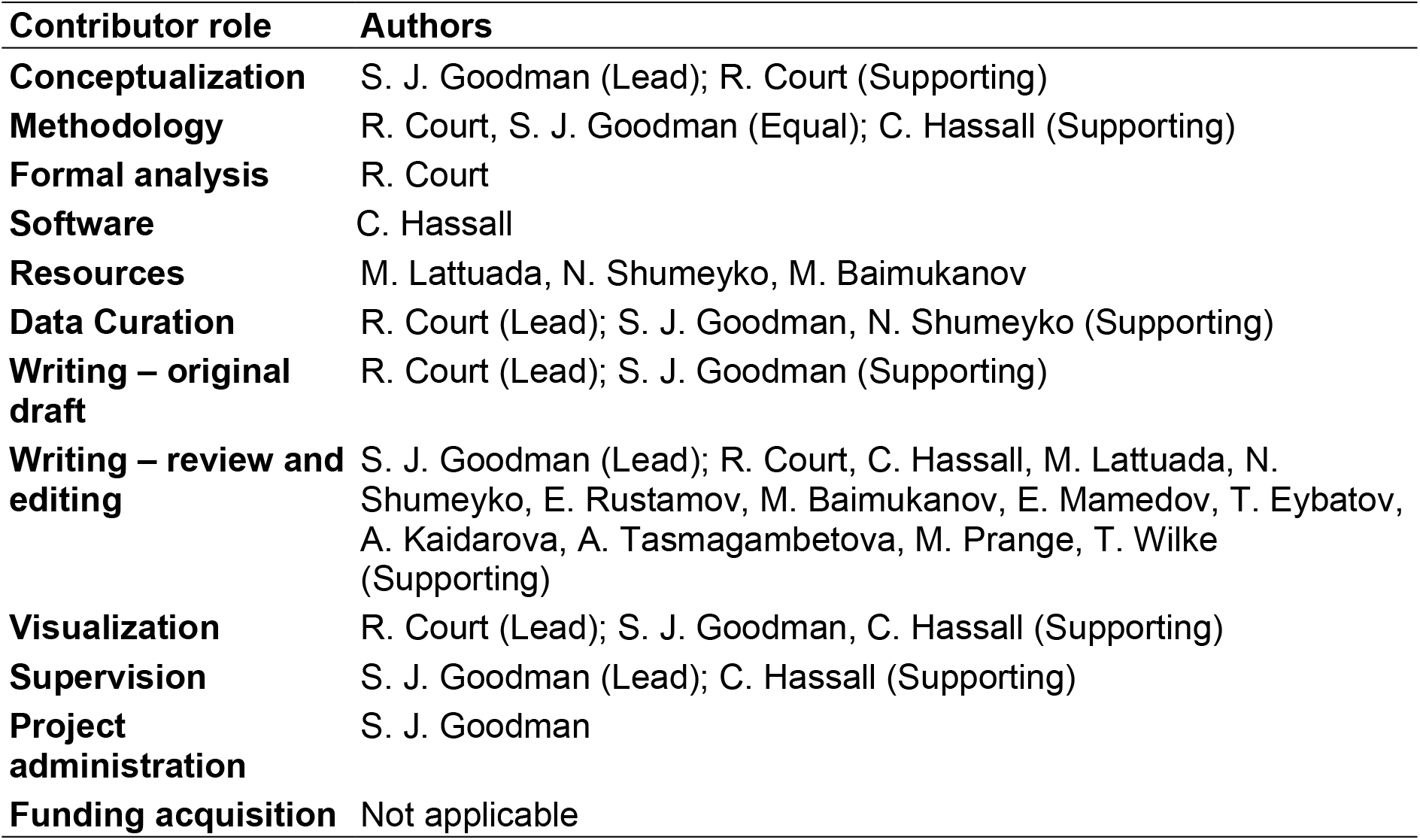

| Contributor role | Authors |
| --- | --- |
| **Conceptualization** | S. J. Goodman (Lead); R. Court (Supporting) |
| **Methodology** | R. Court, S. J. Goodman (Equal); C. Hassall (Supporting) |
| **Formal analysis** | R. Court |
| **Software** | C. Hassall |
| **Resources** | M. Lattuada, N. Shumeyko, M. Baimukanov |
| **Data Curation** | R. Court (Lead); S. J. Goodman, N. Shumeyko (Supporting) |
| **Writing – original draft** | R. Court (Lead); S. J. Goodman (Supporting) |
| **Writing – review and editing** | S. J. Goodman (Lead); R. Court, C. Hassall, M. Lattuada, N. Shumeyko, E. Rustamov, M. Baimukanov, E. Mamedov, T. Eybatov, A. Kaidarova, A. Tasmagambetova, M. Prange, T. Wilke (Supporting) |
| **Visualization** | R. Court (Lead); S. J. Goodman, C. Hassall (Supporting) |
| **Supervision** | S. J. Goodman (Lead); C. Hassall (Supporting) |
| **Project administration** | S. J. Goodman |
| **Funding acquisition** | Not applicable |

## Supplementary Figures and Tables

**Supplementary Table 1**. Ecoregions of the Caspian Sea.

**Supplementary Table 2**. Curatorial corrections/changes made to Protected Area data.

**Supplementary Figure 1**. Caspian Sea bathymetry decline scenarios.

**Supplementary Table 3**. Proportional reduction in water coverage of Important Ecologically or Biologically Significant Areas (EBSAs)

**Supplementary Table 4**. Proportional reduction in water coverage of Important Marine Mammal Areas (IMMAs)

**Supplementary Table 5**. Proportional reduction in water coverage of Caspian Sea ecoregions

**Supplementary Table 6**. Proportional reduction in water coverage of seasonal sturgeonranges

**Supplementary Table 7**. Proportional reduction in water coverage of Marine Protected Areas

**Supplementary Table 8**. Change in distance to shore for coastal and terrestrial protected areas

**Supplementary Table 9**. Change in distance to shore for human settlements

**Supplementary Table 10**. Change in distance to shore for infrastructure

## Data availability

All data used in this study were derived from publicly available sources as specified in the text. GIS and .kml files for decline scenarios, spatial conservation designations, and human infrastructure, along with text files of numerical results and image files are available from: xxxxxxxxxxxxxxxx. A Shiny app to visualise the spatial data is available: xxxxxxx

