## Supplementary material for "Rapid Caspian sea level decline requires dynamic spatio-temporal conservation planning to keep biodiversity protection measures relevant": Court_et_al_2024_Sup_info_part_1

*** Correspondence:**

| | |
|---|---|
| **Supplementary Table 1.** | Ecoregions of the Caspian Sea. |
| **Supplementary Table 2.** | Curatorial corrections/changes made to Protected Area data. |
| **Supplementary Figure 1.** | Caspian Sea bathymetry decline scenarios. |
| **Supplementary Table 3.** | Proportional reduction in water coverage of Important Ecologically or Biologically Significant Areas (EBSAs) |
| **Supplementary Table 4.** | Proportional reduction in water coverage of Important Marine Mammal Areas (IMMAs) |
| **Supplementary Table 5.** | Proportional reduction in water coverage of Caspian Sea ecoregions |
| **Supplementary Table 6.** | Proportional reduction in water coverage of seasonal sturgeon ranges |
| **Supplementary Table 7.** | Proportional reduction in water coverage of Marine Protected Areas |
| **Supplementary Table 8.** | Change in distance to shore for coastal and terrestrial protected areas |

### Definition of ecoregions of the Caspian Sea

Fendereski *et al*., (2014) produced an ecogeographical classification of the Caspian Sea based on physical information derived from remote sensing and in situ data. The authors used a two-step classification procedure, consisting of (i) a data reduction with self-organizing maps (SOMs) and (ii) a synthesis of the most relevant features into a reduced number of marine ecoregions using the hierarchical agglomerative clustering (HAC) method. Six independent variables were used; sea surface temperature, bathymetry, sea ice, seasonal variation of sea surface salinity, total suspended matter and its seasonal variation. The classification yielded 10 distinct ecoregions, separating the northern and middle/southern basins, as well as differentiating nearshore and offshore waters.

Evaluation of differences in community composition based on recorded presence–absence patterns of 25 different species of plankton, fish and benthic invertebrate validates the ecoregions as proxies for habitats with common biological characteristics.

**Supplementary Table 1.** Ecoregions of the Caspian Sea.

| Ecoregion | Acronym | Description/Characteristics |
|---|---|---|
| Northern Caspian Basin River Outflows | NCB-RO | Shallow, dominated by the Volga River outflows, high in sea surface salinity seasonality, high in total suspended matter and its seasonality, covered in ice in winter. |
| Northern Caspian Basin Western shelf | NCB-WS | Shallow, low in sea surface salinity and high in its seasonality, high in total suspended matter, covered in ice in winter. |
| Northern Caspian Basin Ural Furrow | NCB-UF | Shallow, low in sea surface temperature, low in sea surface salinity and its seasonality, covered in ice in winter. |
| Northern Caspian Basin Easternmost shelf | NCB-ES | Shallow, low in sea surface temperature, low in seasonal variation of sea surface salinity, high in total suspended matter and its seasonality, highly ice covered in winter. |
| Northern Caspian Basin Transition | NCB-T | Shallow, high in sea surface salinity seasonality, partly ice covered in winter. |
| Middle Caspian Basin Transition | MCB-T | Medium sea surface temperature, medium seasonal variation of sea surface salinity, medium total suspended matter and DTSM, partly ice covered in winter. |
| Middle Caspian Basin Coastal | MCB-C | Continental shelf area, High in sea surface salinity and low in seasonal variation of sea surface salinity, low in total suspended matter and its seasonality. |
| Middle Caspian Basin Offshore | MCB-OS | Deep, low in sea surface temperature, low in total suspended matter and its seasonality. |
| Southern Caspian Basin Coastal | SCB-C | Continental shelf area, High in sea surface temperature and sea surface salinity, low in total suspended matter and its seasonality. |
| Southern Caspian Basin Offshore | SCB-OS | Deep, high in sea surface temperature, low in total suspended matter and its seasonality. |

### Curation of Protected Area Data

**Supplementary Table 2.** Curatorial corrections/changes made to Protected Area data.

| **Site Name Changes** | | | |
|---|---|---|---|
| Type | Study number designation | Original Name (World Database on Protected Areas* (WDPA) data) | Corrected Local Name |
| Marine | 3 | State Nature Reservat “Akzhaiyk” (1) | Akzhaiyk State Nature Reserve (A) |
| Marine | 4 | State Nature Reservat “Akzhaiyk” (2) | Akzhaiyk State Nature Reserve (B) |
| Marine | 5 | Bujagh National Park | Boujaq National Park and Protected Area |
| Marine | 7 | Volga Delta | Volga Delta Ramsar Site |
| Marine | 8 | Gil Island State Nature Sanctuary | Gil Adasi State Nature Sanctuary |
| Marine | 9 | National Gizilaghaj Park | Gizilaghaj National Park |
| Marine | 12 | Miankaleh Peninsula, Gorgan Bay and Lapoo-Zaghmarz Ab-bandan | Miankaleh Peninsula, Gorgan Bay and Lapoo-Zaghmarz Ramsar Site |
| Terrestrial | *v* | Karadegishskii | Karadegishskiy State Nature Reserve |
| Terrestrial | *vi* | State Nature Sanctuary Karakiya-Karakol (zoological) | Karakiya-Karakol State Nature Sanctuary |
| Terrestrial | *vii* | Kenderly-Kayasan State Reserved Zone | Kendirli-Kayasan State Reserved Zone |
| Terrestrial | *viii* | Lavandavil | Lavandevil Wildlife Refuge |
| Terrestrial | *x* | Samurskij | Samurskiy National Park (Samur Delta Site) |
| Terrestrial | *xii* | Sulakskaya laguna | Sulakskaya Lagoon Firth-Flooded Complex |

| **Designation changes (marine/terrestrial or coastal)** *The designation of the following marine or coastal protection areas were altered from the WDPA designation to better reflect their relation to the Caspian Sea shoreline* | | | |
|---|---|---|---|
| Study number designation | Name | Change | Justification |
| 9 | Gizilaghaj National Park | Terrestrial to Marine | Area largely overlapped with polygon for the water of the Caspian Sea. The polygons of WDPA’s National Giziliaghaj Park (WDPA ID 555651680) and Lesser Gizilaghaj State |

| | | | |
|---|---|---|---|
| | | | Nature Sanctuary (WDPA ID 94019) were merged |
| 15/*x* | Samurskiy National Park (Samur Delta) | The WDPA data consisted of an inland polygon and shore based/marine polygons. The inland polygon was discarded, and only the shoreline/marine based Samur Delta site was retained, with terrestrial and marine components evaluated separately. | |
| *xii* | Sulakskaya Lagoon Firth-Flooded Complex | Marine to Terrestrial | Sulakskaya Lagoon is not part of the Caspian Sea area, but separated from the sea by a sand bar ~100m wide and up to 2.5 m in high. |

| **Exclusions and additions to WDPA data** |
|---|
| The WDPA polygon for the Astrakhanskiy State Nature Biosphere Reserve was removed from analysis as its entire area was represented within the Volga Delta Ramsar Site |
| Shapefiles for the Dagestanskiy State Nature Biosphere Reserve (Kizlyar Bay site) were provided by Dagestanskiy State Nature Biosphere Reserve office. |
| No GIS shape file was available for the Samur-Yalama National Park, Azerbaijan, a counterpart to the Samurskiy National Park in Russia (15/x). The site is shore-based and 117.7km$^2$, and is contained within the area of the Samur-Yalma EBSA. The distance to shore to increase is represented by that for site x. See https://en.wikipedia.org/wiki/Samur-Yalama_National_Park |
| Malyy Zhemchuzhnyy Island State Nature Monument (Russia, 45°1'34'N 48°18'50'E, 35 ha). Being an island terrestrial site in the North there is no appropriate method to perform an increase in distance to shore calculation consistent with the rest of our study. https://www.keybiodiversityareas.org/site/factsheet/1600 |
| No GIS resource was available for the Adamtas State Nature Reserve (zakaznik), Mangystau Region, Karakiya District, Kazakhstan. This has a marine water area of 1546 ha, which is not significant at the Caspian-wide scale of our study. https://tehranconvention.org/system/files/russia/publication_part-2_eng_1.pdf |
| Any other protected areas in planning, or not officially implemented as of July 2023 were not considered. |

* https://www.protectedplanet.net/en/thematic-areas/wdpa?tab=WDPA

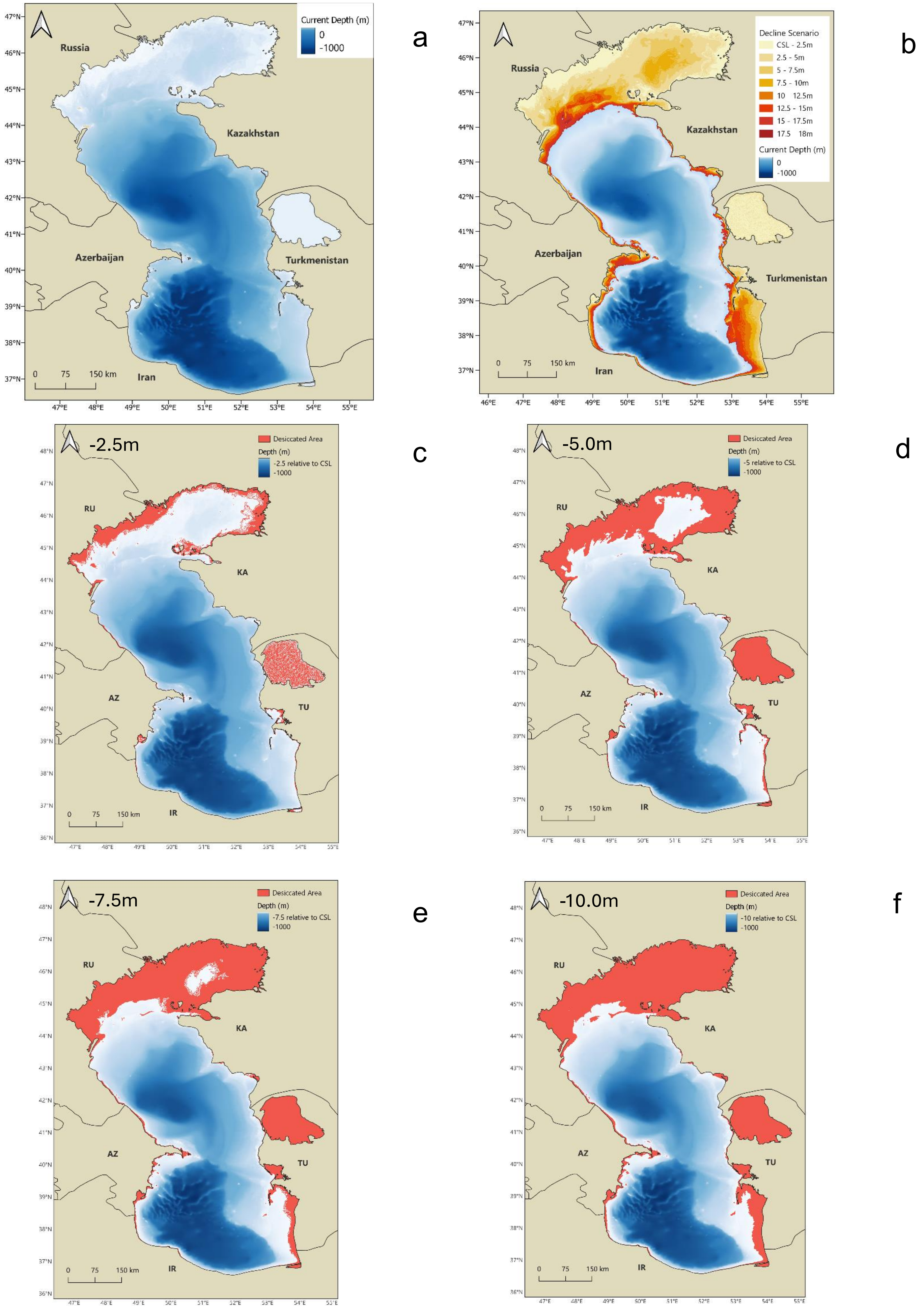

a
Current Depth (m)
0
-1000
Russia
Kazakhstan
Azerbaijan
Turkmenistan
Iran
0 75 150 km
47°N
46°N
45°N
44°N
43°N
42°N
41°N
40°N
39°N
38°N
37°N
47°E 48°E 49°E 50°E 51°E 52°E 53°E 54°E 55°E
b
Decline Scenario
CSL - 2.5m
2.5 - 5m
5 - 7.5m
7.5 - 10m
10 12.5m
12.5 - 15m
15 - 17.5m
17.5 18m
Current Depth (m)
0
-1000
Russia
Kazakhstan
Azerbaijan
Turkmenistan
Iran
0 75 150 km
46°E 47°E 48°E 49°E 50°E 51°E 52°E 53°E 54°E 55°E
c
-2.5m
Desiccated Area
Depth (m)
-2.5 relative to CSL
-1000
RU
KA
AZ
TU
IR
0 75 150 km
d
-5.0m
Desiccated Area
Depth (m)
-5 relative to CSL
-1000
RU
KA
AZ
TU
IR
0 75 150 km
e
-7.5m
Desiccated Area
Depth (m)
-7.5 relative to CSL
-1000
RU
KA
AZ
TU
IR
0 75 150 km
f
-10.0m
Desiccated Area
Depth (m)
-10 relative to CSL
-1000
RU
KA
AZ
TU
IR
0 75 150 km

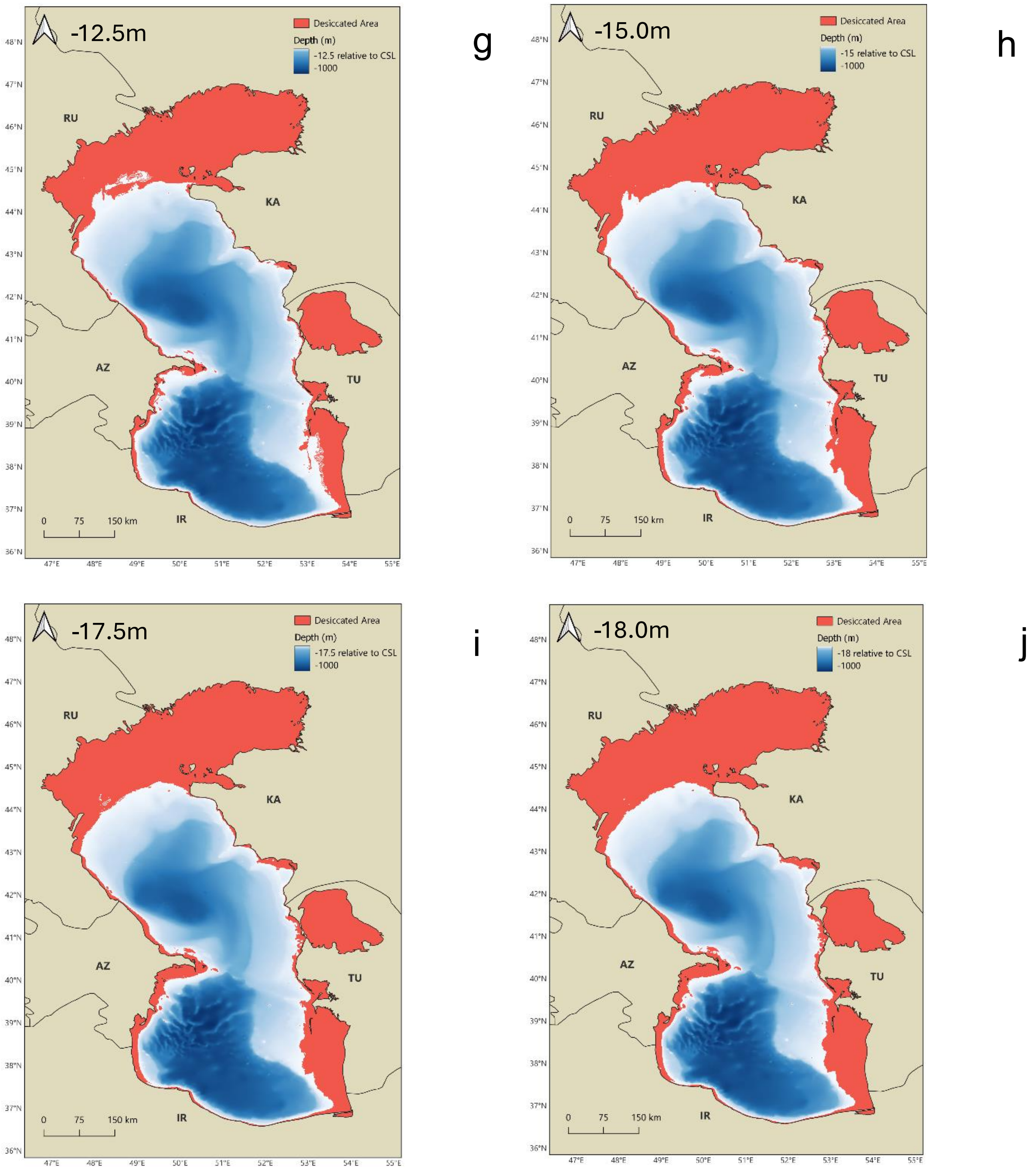


**Supplementary Figure 1.** Caspian Sea bathymetry decline scenarios.

The Caspian Sea current bathymetry (a), and projected water area reduction under eight sea level decline scenarios (b). The area of water that becomes desiccated is shown in red under each individual decline scenario; 2.5m (c), 5m (d), 7.5m (e), 10m (f), 12.5m (g), 15m (h), 17.5m (i), and 18m (j).

**Supplementary Table 3.** The proportional reduction in water coverage of Important Ecologically or Biologically Significant Areas (EBSAs) in the Caspian Sea under different sea level decline scenarios.

| | Sea Level at -27.5m datum | | Decline Scenario | | | | | | | | | | | | | | | |
|---|---|---|---|---|---|---|---|---|---|---|---|---|---|---|---|---|---|---|
| | | | 2.5m | | 5m | | 7.5m | | 10m | | 12.5m | | 15m | | 17.5m | | 18m | |
| EBSA | Total Area km$^2$ | Area intersecting the Caspian Sea, km$^2$ | Area km$^2$ | % Proportion of original area | Area km$^2$ | % Proportion of original area | Area km$^2$ | % Proportion of original area | Area km$^2$ | % Proportion of original area | Area km$^2$ | % Proportion of original area | Area km$^2$ | % Proportion of original area | Area km$^2$ | % Proportion of original area | Area km$^2$ | % Proportion of original area |
| Anzali Wetlands Complex | 248.44 | 0.16 | 0.00 | 0.00 | 0.00 | 0.00 | 0.00 | 0.00 | 0.00 | 0.00 | 0.00 | 0.00 | 0.00 | 0.00 | 0.00 | 0.00 | 0.00 | 0.00 |
| Gizilagach Bay Complex | 2526.16 | 1673.93 | 1277.60 | 76.32 | 1078.77 | 64.45 | 974.00 | 58.19 | 849.75 | 50.76 | 726.00 | 43.37 | 634.92 | 37.93 | 514.64 | 30.74 | 495.66 | 29.61 |
| Karabogazgol Strait | 4111.14 | 2929.83 | 1669.70 | 56.99 | 1158.50 | 39.54 | 1098.90 | 37.51 | 1030.10 | 35.16 | 923.60 | 31.52 | 770.10 | 26.28 | 404.60 | 13.81 | 336.80 | 11.50 |
| Kendirli Bay | 328.78 | 262.52 | 234.80 | 89.44 | 120.45 | 45.88 | 49.83 | 18.98 | 34.49 | 13.14 | 20.96 | 7.98 | 8.58 | 3.27 | 0.00 | 0.00 | 0.00 | 0.00 |
| Kizlyar Bay | 9953.87 | 6607.26 | 3927.66 | 59.44 | 926.31 | 14.02 | 304.76 | 4.61 | 105.11 | 1.59 | 7.76 | 0.12 | 0.00 | 0.00 | 0.00 | 0.00 | 0.00 | 0.00 |
| Komsomol Bay | 1342.89 | 477.35 | 0.00 | 0.00 | 0.00 | 0.00 | 0.00 | 0.00 | 0.00 | 0.00 | 0.00 | 0.00 | 0.00 | 0.00 | 0.00 | 0.00 | 0.00 | 0.00 |
| Kura Delta | 1006.85 | 725.84 | 687.39 | 94.70 | 613.47 | 84.52 | 554.24 | 76.36 | 503.91 | 69.42 | 439.23 | 60.51 | 377.69 | 52.03 | 325.71 | 44.87 | 314.49 | 43.33 |
| Miankaleh-Esenguly | 6409.49 | 5181.83 | 4846.22 | 93.52 | 4122.36 | 79.55 | 3026.43 | 58.40 | 1839.92 | 35.51 | 1397.88 | 26.98 | 1048.58 | 20.24 | 784.91 | 15.15 | 739.86 | 14.28 |
| Pre-estuarine Area of the Ural River | 3209.81 | 1959.54 | 516.29 | 26.35 | 0.00 | 0.00 | 0.00 | 0.00 | 0.00 | 0.00 | 0.00 | 0.00 | 0.00 | 0.00 | 0.00 | 0.00 | 0.00 | 0.00 |
| Pre-estuarine Area of the Volga River | 15232.13 | 11816.81 | 1936.94 | 16.39 | 0.00 | 0.00 | 0.00 | 0.00 | 0.00 | 0.00 | 0.00 | 0.00 | 0.00 | 0.00 | 0.00 | 0.00 | 0.00 | 0.00 |
| Samur-Yalama | 1247.67 | 1066.89 | 1044.62 | 97.91 | 1007.00 | 94.39 | 959.48 | 89.93 | 922.19 | 86.44 | 889.68 | 83.39 | 856.68 | 80.30 | 822.20 | 77.06 | 814.77 | 76.37 |
| Caspian Seal Breeding Grounds | 30891.84 | 30891.84 | 29649.83 | 95.98 | 20167.78 | 65.29 | 12167.27 | 39.39 | 5591.36 | 18.10 | 2618.39 | 8.48 | 826.32 | 2.67 | 535.92 | 1.73 | 479.16 | 1.55 |
| Sefidroud Delta | 139.98 | 117.81 | 110.06 | 93.42 | 101.97 | 86.55 | 93.72 | 79.55 | 85.47 | 72.55 | 76.40 | 64.85 | 62.37 | 52.94 | 43.07 | 36.55 | 39.60 | 33.61 |
| Turkmen Aylagy | 3703.62 | 3384.81 | 3158.43 | 93.31 | 2716.40 | 80.25 | 1699.83 | 50.22 | 647.96 | 19.14 | 0.17 | 0.00 | 0.00 | 0.00 | 0.00 | 0.00 | 0.00 | 0.00 |
| Turkmenbashi Gulf | 2138.15 | 1524.11 | 1010.96 | 66.33 | 260.54 | 17.09 | 22.77 | 1.49 | 0.50 | 0.03 | 0.00 | 0.00 | 0.00 | 0.00 | 0.00 | 0.00 | 0.00 | 0.00 |

**Supplementary Table 4.** The proportional reduction in water coverage of Important Marine Mammal Areas (IMMAs) in the Caspian Sea under different sea level decline scenarios.

| | **Sea Level at -27.5m datum** | | **Decline Scenario** | | | | | | | | | | | | | | | |
|---|---|---|---|---|---|---|---|---|---|---|---|---|---|---|---|---|---|---|
| | | | **2.5m** | | **5m** | | **7.5m** | | **10m** | | **12.5m** | | **15m** | | **17.5m** | | **18m** | |
| IMMA | Total Area km$^2$ | Area intersecting the Caspian Sea km$^2$ | Area km$^2$ | % Proportion of original area | Area km$^2$ | % Proportion of original area | Area km$^2$ | % Proportion of original area | Area km$^2$ | % Proportion of original area | Area km$^2$ | % Proportion of original area | Area km$^2$ | % Proportion of original area | Area km$^2$ | % Proportion of original area | Area km$^2$ | % Proportion of original area |
| Caspian Seal Breeding Area | 55864.36 | 55864.36 | 46437.30 | 83.13 | 23793.00 | 42.59 | 11604.60 | 20.77 | 5078.40 | 9.09 | 2339.55 | 4.19 | 711.60 | 1.27 | 451.20 | 0.81 | 401.70 | 0.72 |
| Caspian Seal Moulting and Haul Out Areas | 27939.24 | 27939.24 | 18228.90 | 65.24 | 9515.40 | 34.06 | 6054.90 | 21.67 | 3629.70 | 12.99 | 2275.20 | 8.14 | 1746.00 | 6.25 | 1108.35 | 3.97 | 994.20 | 3.56 |
| Caspian Seal Transitory Migration and Feeding Area | 163646.05 | 163646.05 | 151846.95 | 92.79 | 141271.80 | 86.33 | 129990.00 | 79.43 | 122078.55 | 74.60 | 116679.75 | 71.30 | 111857.40 | 68.35 | 106974.90 | 65.37 | 106045.05 | 64.80 |

**Supplementary Table 5.** The proportional reduction in water coverage of ecoregions in the Caspian Sea under different sea level decline scenarios.

| | | | Decline Scenario | | | | | | | | | | | | | | | |
|---|---|---|---|---|---|---|---|---|---|---|---|---|---|---|---|---|---|---|
| | | Sea Level at -27.5m datum | 2.5m | | 5m | | 7.5m | | 10m | | 12.5m | | 15m | | 17.5m | | 18m | |
| Ecoregion | Shorthand | Total Area km$^2$ | Area km$^2$ | % proportion of original area | Area km$^2$ | % proportion of original area | Area km$^2$ | % proportion of original area | Area km$^2$ | % proportion of original area | Area km$^2$ | % proportion of original area | Area km$^2$ | % proportion of original area | Area km$^2$ | % proportion of original area | Area km$^2$ | % proportion of original area |
| Middle Caspian Basin - Coastal | mcb-c | 69551.19 | 69348.48 | 99.71 | 68695.77 | 98.77 | 67789.06 | 97.47 | 66533.03 | 95.66 | 64754.21 | 93.10 | 62721.37 | 90.18 | 60311.16 | 86.71 | 59869.49 | 86.08 |
| Middle Caspian Basin - Offshore | mcb-os | 55633.34 | 55603.86 | 99.95 | 55543.04 | 99.84 | 55444.38 | 99.66 | 55385.36 | 99.55 | 55339.06 | 99.47 | 55296.32 | 99.39 | 55252.62 | 99.32 | 55243.78 | 99.30 |
| Middle Caspian Basin - Transition | mcb-t | 30619.52 | 29405.90 | 96.04 | 26669.39 | 87.10 | 25003.20 | 81.66 | 23638.28 | 77.20 | 22591.67 | 73.78 | 21600.03 | 70.54 | 20601.00 | 67.28 | 20410.19 | 66.66 |
| Northern Caspian Basin - Easternmost Shelf | ncb-es | 11359.10 | 4962.38 | 43.69 | 68.05 | 0.60 | 0.00 | 0.00 | 0.00 | 0.00 | 0.00 | 0.00 | 0.00 | 0.00 | 0.00 | 0.00 | 0.00 | 0.00 |
| Northern Caspian Basin - River Outflows | ncb-ro | 8385.90 | 2839.72 | 33.86 | 495.46 | 5.91 | 4.60 | 0.05 | 0.00 | 0.00 | 0.00 | 0.00 | 0.00 | 0.00 | 0.00 | 0.00 | 0.00 | 0.00 |
| Northern Caspian Basin - Transition | ncb-t | 21941.55 | 21522.54 | 98.09 | 16303.38 | 74.30 | 12178.06 | 55.50 | 9474.71 | 43.18 | 5930.38 | 27.03 | 3464.93 | 15.79 | 1711.82 | 7.80 | 1444.20 | 6.58 |
| Northern Caspian Basin - Ural Furrow | ncb-uf | 17094.46 | 15782.79 | 92.33 | 10720.80 | 62.72 | 3951.46 | 23.12 | 0.00 | 0.00 | 0.00 | 0.00 | 0.00 | 0.00 | 0.00 | 0.00 | 0.00 | 0.00 |
| Northern Caspian Basin - Western Shelf | ncb-ws | 18963.32 | 12957.72 | 68.33 | 2765.90 | 14.59 | 357.88 | 1.89 | 0.00 | 0.00 | 0.00 | 0.00 | 0.00 | 0.00 | 0.00 | 0.00 | 0.00 | 0.00 |
| Southern Caspian Basin - Coastal | scb-c | 52461.16 | 51394.57 | 97.97 | 49126.38 | 93.64 | 44949.70 | 85.68 | 40676.57 | 77.54 | 35459.33 | 67.59 | 30979.63 | 59.05 | 28495.96 | 54.32 | 28085.77 | 53.54 |
| Southern Caspian Basin - Offshore | scb-os | 80336.73 | 80328.88 | 99.99 | 80299.82 | 99.95 | 80250.18 | 99.89 | 80183.46 | 99.81 | 80092.49 | 99.70 | 79991.83 | 99.57 | 79870.21 | 99.42 | 79843.56 | 99.39 |

**Supplementary Table 6.** The proportional reduction in water coverage of seasonal sturgeon ranges in the Caspian Sea under different sea level decline scenarios.

| | | **Decline Scenario** | | | | | | | | | | | | | | | |
|---|---|---|---|---|---|---|---|---|---|---|---|---|---|---|---|---|---|
| | **Sea Level at -27.5m datum** | **2.5m** | | **5m** | | **7.5m** | | **10m** | | **12.5m** | | **15m** | | **17.5m** | | **18m** | |
| Season | Total Area $km^2$ | Area $km^2$ | % proportion of original area | Area $km^2$ | % proportion of original area | Area $km^2$ | % proportion of original area | Area $km^2$ | % proportion of original area | Area $km^2$ | % proportion of original area | Area $km^2$ | % proportion of original area | Area $km^2$ | % proportion of original area | Area $km^2$ | % proportion of original area |
| Summer/ Autumn | 141149.20 | 132972.17 | 94.21 | 107030.10 | 75.83 | 89549.30 | 63.44 | 78689.95 | 55.75 | 70337.23 | 49.83 | 62754.85 | 44.46 | 57662.31 | 40.85 | 56800.24 | 40.24 |
| Winter/Spring | 103167.34 | 102998.79 | 99.84 | 102617.16 | 99.47 | 101411.59 | 98.30 | 98560.00 | 95.53 | 94183.74 | 91.29 | 89792.87 | 87.04 | 86955.94 | 84.29 | 86459.84 | 83.81 |

**Supplementary Table 7.** The proportional reduction in water coverage of Marine Protected Areas in the Caspian Sea under different sea level decline scenarios.

| | | Sea Level at -27.5m datum | | Decline Scenario | | | | | | | | | | | | | | | |
|---|---|---|---|---|---|---|---|---|---|---|---|---|---|---|---|---|---|---|---|
| | | | | 2.5m | | 5m | | 7.5m | | 10m | | 12.5m | | 15m | | 17.5m | | 18m | |
| Protected Area | Category/ Description | Total Area, km$^2$ | Area intersecting the Caspian Sea, km$^2$ | Area km$^2$ | % Proportion of original area | Area km$^2$ | % Proportion of original area | Area km$^2$ | % Proportion of original area | Area km$^2$ | % Proportion of original area | Area km$^2$ | % Proportion of original area | Area km$^2$ | % Proportion of original area | Area km$^2$ | % Proportion of original area | Area km$^2$ | % Proportion of original area |
| Gizilaghaj National Park | National Park | 1083.57 | 424.99 | 72.60 | 17.08 | 0.00 | 0.00 | 0.00 | 0.00 | 0.00 | 0.00 | 0.00 | 0.00 | 0.00 | 0.00 | 0.00 | 0.00 | 0.00 | 0.00 |
| Absheron National Park | National Park | 7.85 | 6.32 | 0.14 | 2.15 | 0.00 | 0.00 | 0.00 | 0.00 | 0.00 | 0.00 | 0.00 | 0.00 | 0.00 | 0.00 | 0.00 | 0.00 | 0.00 | 0.00 |
| Agrahanskiy State Nature Sanctuary | State Nature Sanctuary | 406.52 | 147.56 | 2.40 | 1.63 | 0.00 | 0.00 | 0.00 | 0.00 | 0.00 | 0.00 | 0.00 | 0.00 | 0.00 | 0.00 | 0.00 | 0.00 | 0.00 | 0.00 |
| Akzhaiyk State Nature Reserve (A) | State Nature Reserve | 517.34 | 351.96 | 28.80 | 8.18 | 0.00 | 0.00 | 0.00 | 0.00 | 0.00 | 0.00 | 0.00 | 0.00 | 0.00 | 0.00 | 0.00 | 0.00 | 0.00 | 0.00 |
| Akzhaiyk State Nature Reserve (B) | State Nature Reserve | 597.59 | 254.45 | 24.00 | 9.43 | 0.00 | 0.00 | 0.00 | 0.00 | 0.00 | 0.00 | 0.00 | 0.00 | 0.00 | 0.00 | 0.00 | 0.00 | 0.00 | 0.00 |
| Boujaq National Park | Ramsar Site, Wetland of International Importance | 32.69 | 20.35 | 9.60 | 47.18 | 4.20 | 20.64 | 1.05 | 5.16 | 0.09 | 0.43 | 0.00 | 0.00 | 0.00 | 0.00 | 0.00 | 0.00 | 0.00 | 0.00 |
| Gil Adasi State Nature Sanctuary | State Nature Sanctuary | 1.53 | 1.45 | 0.00 | 0.00 | 0.00 | 0.00 | 0.00 | 0.00 | 0.00 | 0.00 | 0.00 | 0.00 | 0.00 | 0.00 | 0.00 | 0.00 | 0.00 | 0.00 |
| Hazar State Nature Reserve (A) | State Nature Reserve | 669.93 | 337.51 | 188.70 | 55.91 | 55.20 | 16.36 | 0.00 | 0.00 | 0.00 | 0.00 | 0.00 | 0.00 | 0.00 | 0.00 | 0.00 | 0.00 | 0.00 | 0.00 |
| Hazar State Nature Reserve (B) | State Nature Reserve | 1958.27 | 1605.49 | 926.70 | 57.72 | 238.35 | 14.85 | 23.25 | 1.45 | 0.90 | 0.06 | 0.00 | 0.00 | 0.00 | 0.00 | 0.00 | 0.00 | 0.00 | 0.00 |

| | | | | | | | | | | | | | | | | | | | |
|---|---|---|---|---|---|---|---|---|---|---|---|---|---|---|---|---|---|---|---|
| Miankaleh Protected Area | UNESCO-MAB Biosphere Reserve | 708.85 | 389.98 | 154.20 | 39.54 | 0.75 | 0.19 | 0.00 | 0.00 | 0.00 | 0.00 | 0.00 | 0.00 | 0.00 | 0.00 | 0.00 | 0.00 | 0.00 | 0.00 |
| Miankaleh Peninsula, Gorgan Bay and Lapoo-Zaghmarz Ramsar Site | Ramsar Site, Wetland of International Importance | 959.09 | 361.94 | 147.00 | 40.61 | 0.00 | 0.00 | 0.00 | 0.00 | 0.00 | 0.00 | 0.00 | 0.00 | 0.00 | 0.00 | 0.00 | 0.00 | 0.00 | 0.00 |
| Novinskiy State Nature Sanctuary | State Nature Sanctuary | 494.14 | 342.16 | 0.00 | 0.00 | 0.00 | 0.00 | 0.00 | 0.00 | 0.00 | 0.00 | 0.00 | 0.00 | 0.00 | 0.00 | 0.00 | 0.00 | 0.00 | 0.00 |
| Volga Delta Ramsar Site | Ramsar Site, Wetland of International Importance | 10646.97 | 7035.50 | 119.70 | 1.70 | 0.00 | 0.00 | 0.00 | 0.00 | 0.00 | 0.00 | 0.00 | 0.00 | 0.00 | 0.00 | 0.00 | 0.00 | 0.00 | 0.00 |
| State Reserved Zone in Northern part of Caspian Sea | State Natural Protected Zone | 63052.14 | 56643.74 | 43121.85 | 76.13 | 20917.2 | 36.93 | 8922.0 | 15.75 | 2970.15 | 5.24 | 2314.35 | 4.09 | 1883.25 | 3.32 | 1533.00 | 2.71 | 1494.3 | 2.64 |
| Samurskiy National Park (Samur Delta) | National Park | 121.60 | 33.20 | 29.20 | 87.95 | 23.52 | 70.82 | 17.51 | 52.74 | 11.60 | 34.93 | 6.68 | 20.11 | 3.07 | 9.26 | 0.55 | 1.66 | 0.28 | 0.84 |
| Dagestanskiy State Nature Biosphere Reserve (Kizlyar Bay) | State Nature Biosphere Reserve | 383.75 | 294.47 | 284.22 | 96.52 | 1.23 | 0.42 | 0.00 | 0.00 | 0.00 | 0.00 | 0.00 | 0.00 | 0.00 | 0.00 | 0.00 | 0.00 | 0.00 | 0.00 |

**Supplementary Table 8.** The change in distance to shore for coastal and terrestrial protected areas, from their centroid, in the Caspian Sea under different sea level decline scenarios.

| | | Sea Level at -27.5m datum | Decline Scenario | | | | | | | | | | | | | | | |
|---|---|---|---|---|---|---|---|---|---|---|---|---|---|---|---|---|---|---|
| | | | 2.5m | | 5m | | 7.5m | | 10m | | 12.5m | | 15m | | 17.5m | | 18m | |
| Protected Area | Category/ Description | Distance to shore from centroid, km | Distance to Shore, km | Change in distance to shore, km | Distance to Shore, km | Change in distance to shore, km | Distance to Shore, km | Change in distance to shore, km | Distance to Shore, km | Change in distance to shore, km | Distance to Shore, km | Change in distance to shore, km | Distance to Shore, km | Change in distance to shore, km | Distance to Shore, km | Change in distance to shore, km | Distance to Shore, km | Change in distance to shore, km |
| Aktay-Buzachi State Nature Sanctuary | State Nature Sanctuary | 7.22 | 8.68 | 1.46 | 17.24 | 10.02 | 21.10 | 13.88 | 46.02 | 38.79 | 53.36 | 46.14 | 68.28 | 61.05 | 68.71 | 61.49 | 68.93 | 61.71 |
| Alborz-e-Markazy Protected Area | Protected Area | 47.15 | 47.15 | 0.00 | 47.15 | 0.00 | 47.15 | 0.00 | 47.15 | 0.00 | 47.15 | 0.00 | 47.15 | 0.00 | 47.15 | 0.00 | 47.15 | 0.00 |
| Amirkelayeh Lake Ramsar Site | Ramsar Site, Wetland of International Importance | 1.65 | 2.25 | 0.60 | 3.11 | 1.47 | 3.50 | 1.85 | 4.23 | 2.59 | 5.43 | 3.79 | 6.16 | 4.51 | 6.53 | 4.88 | 6.53 | 4.88 |
| Gomishan Lagoon Ramsar Site | Ramsar Site, Wetland of International Importance | 3.15 | 3.15 | 0.00 | 4.50 | 1.35 | 10.48 | 7.33 | 16.24 | 13.09 | 19.46 | 16.31 | 28.92 | 25.76 | 34.32 | 31.17 | 36.00 | 32.85 |
| Karadegishskiy State Nature Reserve | State Nature Reserve | 28.03 | 28.20 | 0.17 | 32.96 | 4.93 | 38.95 | 10.92 | 45.75 | 17.72 | 48.64 | 20.61 | 55.70 | 27.67 | 61.67 | 33.64 | 62.87 | 34.84 |
| Kendirli-Kayasan State Nature Reserved Zone | State Natural Protected Zone | 62.26 | 62.26 | 0.00 | 62.47 | 0.21 | 68.02 | 5.76 | 71.71 | 9.45 | 72.19 | 9.94 | 72.84 | 10.58 | 74.60 | 12.34 | 75.25 | 12.99 |
| Lavandevil Wildlife Refuge | Wildlife Refuge | 0.00 | 0.99 | 0.99 | 2.75 | 2.75 | 4.57 | 4.57 | 5.73 | 5.73 | 7.85 | 7.85 | 12.22 | 12.22 | 13.31 | 13.31 | 13.58 | 13.58 |
| Lisar Protected Area | Protected Area | 14.46 | 14.71 | 0.25 | 15.42 | 0.97 | 16.41 | 1.96 | 17.58 | 3.12 | 19.52 | 5.07 | 19.52 | 5.07 | 19.52 | 5.07 | 19.52 | 5.07 |
| Samurskiy National Park (Samur Delta Site) | National Park | 0.45 | 4.44 | 3.99 | 4.87 | 4.41 | 5.19 | 4.74 | 5.69 | 5.24 | 6.11 | 5.66 | 6.54 | 6.09 | 7.10 | 6.64 | 7.24 | 6.79 |
| Shirvan National Park | National Park | 9.73 | 9.73 | 0.00 | 10.41 | 0.68 | 12.37 | 2.64 | 14.36 | 4.63 | 14.36 | 4.63 | 14.76 | 5.03 | 14.94 | 5.21 | 15.17 | 5.43 |
| Karakiya-Karakol State Nature Sanctuary | State Nature Sanctuary | 1.96 | 2.21 | 0.25 | 2.55 | 0.59 | 2.89 | 0.94 | 3.56 | 1.60 | 4.23 | 2.27 | 5.80 | 3.85 | 7.55 | 5.59 | 8.27 | 6.31 |
| Sulakskaya Lagoon Firth-Flooded Complex | Firth-Flooded Complex | 0.89 | 1.55 | 0.66 | 2.28 | 1.39 | 2.85 | 1.96 | 3.66 | 2.77 | 8.01 | 7.12 | 12.54 | 11.65 | 15.27 | 14.38 | 18.42 | 17.53 |

Intentionally left blank.
