## Supplementary material for "Rapid Caspian sea level decline requires dynamic spatio-temporal conservation planning to keep biodiversity protection measures relevant": Court_et_al_2024_Sup_info_part_2

*** Correspondence:**

**Supplementary Table 9.** Change in distance to shore for coastal settlements

**Supplementary Table 10.** Change in distance to shore for infrastructure

Supplementary Table 9. The change in distance to shore for human settlements, located on the coast of the Caspian Sea, under different sea level decline scenarios.

| | | | | | Decline Scenario | | | | | | | | | | | | | | | |
|---|---|---|---|---|---|---|---|---|---|---|---|---|---|---|---|---|---|---|---|---|
| | | | | Sea level at -27.5m datum | 2.5m | | 5m | | 7.5m | | 10m | | 12.5m | | 15m | | 17.5m | | 18m | |
| Human Settlement | Country | Latitude | Longitude | Distance to shore, km | Distance to shore, km | Change in distance to shore, km | Distance to shore, km | Change in distance to shore, km | Distance to shore, km | Change in distance to shore, km | Distance to shore, km | Change in distance to shore, km | Distance to shore, km | Change in distance to shore, km | Distance to shore, km | Change in distance to shore, km | Distance to shore, km | Change in distance to shore, km | Distance to shore, km | Change in distance to shore, km |
| Neka | Iran | 36.65 | 53.30 | 21.32 | 21.32 | 0.00 | 21.83 | 0.51 | 22.45 | 1.14 | 23.20 | 1.88 | 24.88 | 3.56 | 26.36 | 5.04 | 27.95 | 6.63 | 28.29 | 6.97 |
| Rud Sar | Iran | 37.15 | 50.28 | 2.24 | 2.49 | 0.25 | 2.92 | 0.68 | 3.78 | 1.54 | 4.64 | 2.41 | 5.68 | 3.44 | 6.55 | 4.31 | 7.42 | 5.18 | 7.72 | 5.48 |
| Now Shahr | Iran | 36.60 | 51.50 | 5.48 | 5.48 | 0.00 | 5.70 | 0.22 | 5.84 | 0.36 | 6.12 | 0.64 | 6.40 | 0.92 | 7.27 | 1.80 | 7.96 | 2.49 | 8.11 | 2.63 |
| Astara | Iran | 38.39 | 48.85 | 2.18 | 3.08 | 0.91 | 4.50 | 2.33 | 6.50 | 4.33 | 7.78 | 5.60 | 9.23 | 7.06 | 12.84 | 10.67 | 14.46 | 12.29 | 14.62 | 12.44 |
| Tonekabon | Iran | 36.84 | 50.84 | 0.88 | 0.88 | 0.00 | 0.88 | 0.00 | 1.11 | 0.23 | 1.24 | 0.36 | 1.66 | 0.78 | 2.03 | 1.15 | 2.42 | 1.53 | 2.42 | 1.53 |
| Fereydunkenar | Iran | 36.69 | 52.56 | 0.81 | 0.99 | 0.18 | 1.12 | 0.31 | 1.45 | 0.64 | 1.91 | 1.10 | 2.37 | 1.56 | 2.84 | 2.03 | 3.30 | 2.49 | 3.34 | 2.53 |
| Neftcala | Azerbaijan | 39.37 | 49.26 | 5.32 | 5.62 | 0.30 | 9.21 | 3.88 | 9.48 | 4.15 | 9.78 | 4.46 | 10.05 | 4.73 | 10.32 | 5.00 | 10.62 | 5.30 | 10.62 | 5.30 |
| Fort Shevchenko | Kazakhstan - North basin | 44.51 | 50.26 | 1.80 | 1.86 | 0.06 | 1.86 | 0.06 | 1.86 | 0.06 | 2.52 | 0.72 | 6.47 | 4.67 | 8.69 | 6.89 | 9.98 | 8.18 | 10.24 | 8.43 |
| Zyrya | Azerbaijan | 40.37 | 50.28 | 4.34 | 4.34 | 0.00 | 4.34 | 0.00 | 4.71 | 0.37 | 6.87 | 2.54 | 11.87 | 7.53 | 14.20 | 9.87 | 15.51 | 11.18 | 15.85 | 11.52 |
| Mardakyan | Azerbaijan | 40.49 | 50.16 | 2.32 | 2.70 | 0.38 | 3.11 | 0.78 | 3.47 | 1.15 | 4.11 | 1.79 | 5.41 | 3.09 | 7.18 | 4.86 | 11.31 | 8.99 | 11.75 | 9.42 |
| Qobustan | Azerbaijan | 40.09 | 49.41 | 0.26 | 1.43 | 1.17 | 5.89 | 5.64 | 7.88 | 7.62 | 8.37 | 8.11 | 10.69 | 10.44 | 18.50 | 18.24 | 24.80 | 24.54 | 27.07 | 26.82 |
| Pirallahi | Azerbaijan | 40.47 | 50.34 | 0.34 | 1.01 | 0.67 | 1.31 | 0.97 | 1.86 | 1.52 | 2.16 | 1.82 | 2.89 | 2.55 | 5.41 | 5.06 | 6.30 | 5.96 | 6.43 | 6.09 |
| Hovsan | Azerbaijan | 40.39 | 50.09 | 2.21 | 2.72 | 0.51 | 3.65 | 1.44 | 4.15 | 1.93 | 5.54 | 3.33 | 13.83 | 11.62 | 19.74 | 17.52 | 22.30 | 20.08 | 22.74 | 20.53 |
| Turkan | Azerbaijan | 40.37 | 50.21 | 1.47 | 1.73 | 0.26 | 1.98 | 0.51 | 2.54 | 1.07 | 5.57 | 4.10 | 14.13 | 12.66 | 18.58 | 17.10 | 20.84 | 19.37 | 21.17 | 19.70 |
| Nardaran | Azerbaijan | 40.56 | 49.98 | 2.79 | 3.47 | 0.68 | 3.65 | 0.86 | 3.92 | 1.13 | 4.20 | 1.41 | 4.63 | 1.85 | 5.08 | 2.29 | 5.97 | 3.18 | 6.42 | 3.63 |
| Bekdash | Turkmenistan | 41.55 | 52.58 | 2.11 | 2.11 | 0.00 | 2.11 | 0.00 | 2.11 | 0.00 | 2.29 | 0.18 | 2.48 | 0.37 | 2.73 | 0.62 | 3.02 | 0.91 | 3.10 | 0.99 |
| Fetisovo | Kazakhstan | 42.77 | 52.65 | 1.60 | 1.78 | 0.18 | 2.80 | 1.20 | 7.57 | 5.97 | 12.49 | 10.89 | 14.19 | 12.59 | 15.76 | 14.16 | 17.72 | 16.12 | 19.11 | 17.51 |
| Kuryk | Kazakhstan | 43.18 | 51.68 | 2.05 | 2.19 | 0.14 | 3.14 | 1.09 | 4.10 | 2.05 | 6.22 | 4.17 | 8.31 | 6.26 | 8.89 | 6.84 | 9.46 | 7.41 | 9.56 | 7.52 |
| Xacmas | Azerbaijan | 41.37 | 48.94 | 8.94 | 9.43 | 0.50 | 10.63 | 1.69 | 11.21 | 2.27 | 11.60 | 2.66 | 15.55 | 6.62 | 16.66 | 7.72 | 17.61 | 8.67 | 17.78 | 8.84 |
| Zarat | Azerbaijan | 40.95 | 49.28 | 0.81 | 1.56 | 0.75 | 1.85 | 1.04 | 2.47 | 1.66 | 4.08 | 3.27 | 5.78 | 4.97 | 7.20 | 6.39 | 8.83 | 8.02 | 9.11 | 8.30 |

(continued)

| | | | | | Decline Scenario | | | | | | | | | | | | | | | |
|---|---|---|---|---|---|---|---|---|---|---|---|---|---|---|---|---|---|---|---|---|
| | | | | Sea level at -27.5m datum | 2.5m | | 5m | | 7.5m | | 10m | | 12.5m | | 15m | | 17.5m | | 18m | |
| Human Settlement | Country | Latitude | Longitude | Distance to shore, km | Distance to shore, km | Change in distance to shore, km | Distance to shore, km | Change in distance to shore, km | Distance to shore, km | Change in distance to shore, km | Distance to shore, km | Change in distance to shore, km | Distance to shore, km | Change in distance to shore, km | Distance to shore, km | Change in distance to shore, km | Distance to shore, km | Change in distance to shore, km | Distance to shore, km | Change in distance to shore, km |
| Nabran | Azerbaijan | 41.76 | 48.69 | 0.38 | 0.73 | 0.36 | 1.00 | 0.63 | 1.00 | 0.63 | 1.54 | 1.16 | 1.88 | 1.51 | 2.46 | 2.09 | 2.95 | 2.57 | 2.95 | 2.57 |
| Muxtedir | Azerbaijan | 41.66 | 48.78 | 0.05 | 1.44 | 1.39 | 2.01 | 1.96 | 2.93 | 2.88 | 3.66 | 3.61 | 4.09 | 4.04 | 4.49 | 4.45 | 4.89 | 4.84 | 5.04 | 5.00 |
| Gilazi | Azerbaijan | 40.87 | 49.34 | 2.26 | 3.18 | 0.92 | 4.04 | 1.78 | 6.53 | 4.27 | 6.88 | 4.63 | 7.11 | 4.85 | 7.69 | 5.43 | 9.21 | 6.95 | 9.49 | 7.23 |
| Bandar-e Torkaman | Iran | 36.90 | 54.07 | 2.09 | 2.77 | 0.68 | 12.06 | 9.97 | 16.30 | 14.21 | 21.30 | 19.21 | 28.75 | 26.66 | 38.21 | 36.13 | 43.57 | 41.48 | 45.64 | 43.55 |
| Xudat | Azerbaijan | 41.63 | 48.68 | 8.63 | 9.50 | 0.87 | 10.06 | 1.43 | 10.62 | 1.99 | 11.48 | 2.85 | 12.05 | 3.42 | 12.55 | 3.92 | 12.87 | 4.24 | 13.11 | 4.48 |
| Karshy | Turkmenistan | 40.72 | 52.86 | 0.59 | 0.90 | 0.31 | 1.22 | 0.63 | 1.78 | 1.19 | 2.62 | 2.03 | 3.30 | 2.71 | 3.65 | 3.06 | 4.36 | 3.77 | 4.71 | 4.12 |
| Bautino | Kazakhstan - North basin | 44.54 | 50.26 | 0.30 | 0.30 | 0.00 | 1.06 | 0.77 | 1.53 | 1.23 | 3.47 | 3.17 | 5.88 | 5.59 | 7.61 | 7.32 | 12.64 | 12.34 | 12.96 | 12.66 |
| Kanga | Kazakhstan - North basin | 44.62 | 50.59 | 1.32 | 1.53 | 0.21 | 1.63 | 0.31 | 1.79 | 0.47 | 2.19 | 0.87 | 3.79 | 2.47 | 16.21 | 14.89 | 19.21 | 17.90 | 19.21 | 17.90 |
| Erkinkala | Kazakhstan - North basin | 47.02 | 51.81 | 16.10 | 22.73 | 6.63 | 50.25 | 34.14 | 73.23 | 57.13 | 276.40 | 260.30 | 280.01 | 263.90 | 284.86 | 268.75 | 285.41 | 269.31 | 285.41 | 269.31 |
| Kyzylbalyk | Kazakhstan - North basin | 46.98 | 51.79 | 12.62 | 18.57 | 5.95 | 45.35 | 32.73 | 68.28 | 55.66 | 271.44 | 258.82 | 275.05 | 262.43 | 279.91 | 267.28 | 280.46 | 267.84 | 280.46 | 267.84 |
| Zhanatalap | Kazakhstan - North basin | 47.00 | 51.82 | 15.37 | 21.57 | 6.19 | 48.33 | 32.95 | 70.99 | 55.61 | 274.28 | 258.91 | 277.91 | 262.54 | 282.78 | 267.41 | 283.34 | 267.96 | 283.34 | 267.96 |
| Kurmangazy | Kazakhstan - North basin | 46.97 | 51.78 | 11.43 | 17.17 | 5.74 | 43.92 | 32.49 | 66.95 | 55.52 | 270.05 | 258.62 | 273.65 | 262.22 | 278.50 | 267.07 | 279.06 | 267.63 | 279.06 | 267.63 |
| Damba | Kazakhstan - North basin | 46.96 | 51.75 | 9.58 | 15.21 | 5.62 | 42.21 | 32.62 | 65.53 | 55.95 | 268.48 | 258.89 | 272.06 | 262.47 | 276.88 | 267.30 | 277.44 | 267.86 | 277.44 | 267.86 |
| Peshnoy | Kazakhstan - North basin | 46.91 | 51.68 | 2.47 | 7.24 | 4.77 | 34.48 | 32.01 | 58.74 | 56.27 | 261.02 | 258.55 | 264.53 | 262.06 | 269.31 | 266.84 | 269.86 | 267.39 | 269.86 | 267.39 |
| Taskala | Kazakhstan - North basin | 47.03 | 51.91 | 20.79 | 29.24 | 8.45 | 54.00 | 33.21 | 76.49 | 55.70 | 280.22 | 259.43 | 283.85 | 263.06 | 288.91 | 268.12 | 289.47 | 268.68 | 289.47 | 268.68 |
| Kraynovka | Russia - North basin | 43.98 | 47.38 | 1.33 | 1.78 | 0.45 | 15.75 | 14.41 | 25.50 | 24.17 | 41.97 | 40.64 | 47.96 | 46.63 | 54.05 | 52.72 | 59.74 | 58.41 | 61.64 | 60.31 |
| Noviy Biryuzyak | Russia - North basin | 43.78 | 47.30 | 16.07 | 19.00 | 2.94 | 27.32 | 11.26 | 30.94 | 14.87 | 34.91 | 18.84 | 40.87 | 24.81 | 47.49 | 31.42 | 53.74 | 37.68 | 55.12 | 39.05 |
| Kallektivizator | Russia - North basin | 43.96 | 47.39 | 0.51 | 1.29 | 0.78 | 15.25 | 14.75 | 24.90 | 24.40 | 41.06 | 40.55 | 47.32 | 46.81 | 52.89 | 52.38 | 57.81 | 57.30 | 59.46 | 58.96 |
| Novoterechnoe | Russia - North basin | 44.02 | 47.35 | 1.07 | 1.45 | 0.38 | 14.72 | 13.65 | 26.67 | 25.60 | 42.72 | 41.64 | 48.90 | 47.83 | 55.20 | 54.13 | 62.76 | 61.69 | 65.13 | 64.05 |
| Noviy Bakhtemir | Russia - North basin | 44.13 | 47.27 | 0.00 | 2.16 | 2.16 | 14.07 | 14.07 | 34.08 | 34.08 | 48.08 | 48.08 | 53.82 | 53.82 | 60.12 | 60.12 | 66.66 | 66.66 | 68.75 | 68.75 |

(continued)

| | | | | | Decline Scenario | | | | | | | | | | | | | | | |
|---|---|---|---|---|---|---|---|---|---|---|---|---|---|---|---|---|---|---|---|---|
| | | | | Sea level at -27.5m datum | 2.5m | | 5m | | 7.5m | | 10m | | 12.5m | | 15m | | 17.5m | | 18m | |
| Human Settlement | Country | Latitude | Longitude | Distance to shore, km | Distance to shore, km | Change in distance to shore, km | Distance to shore, km | Change in distance to shore, km | Distance to shore, km | Change in distance to shore, km | Distance to shore, km | Change in distance to shore, km | Distance to shore, km | Change in distance to shore, km | Distance to shore, km | Change in distance to shore, km | Distance to shore, km | Change in distance to shore, km | Distance to shore, km | Change in distance to shore, km |
| Suyutkino | Russia - North basin | 44.19 | 47.24 | 0.00 | 2.59 | 2.59 | 17.05 | 17.05 | 37.97 | 37.97 | 50.00 | 50.00 | 55.73 | 55.73 | 61.99 | 61.99 | 67.80 | 67.80 | 70.05 | 70.05 |
| Krasniy Rybak | Russia - North basin | 44.18 | 47.25 | 0.00 | 2.42 | 2.42 | 16.43 | 16.43 | 37.31 | 37.31 | 49.65 | 49.65 | 55.34 | 55.34 | 61.64 | 61.64 | 67.52 | 67.52 | 69.75 | 69.75 |
| Noviy Chechen | Russia - North basin | 44.26 | 47.07 | 0.31 | 3.08 | 2.77 | 28.60 | 28.29 | 53.29 | 52.98 | 63.54 | 63.22 | 69.91 | 69.60 | 75.70 | 75.39 | 81.18 | 80.87 | 83.50 | 83.19 |
| Bryanskiy Rybozavod | Russia - North basin | 44.31 | 47.04 | 0.07 | 4.86 | 4.79 | 30.47 | 30.39 | 58.00 | 57.92 | 66.17 | 66.10 | 72.41 | 72.34 | 77.88 | 77.81 | 83.76 | 83.68 | 86.03 | 85.96 |
| Bryansk | Russia - North basin | 44.33 | 46.98 | 4.59 | 7.62 | 3.03 | 35.36 | 30.77 | 61.94 | 57.35 | 71.29 | 66.70 | 77.38 | 72.79 | 80.00 | 75.41 | 88.84 | 84.25 | 91.10 | 86.51 |
| M. Gadzhieva | Russia - North basin | 43.98 | 47.32 | 5.51 | 5.75 | 0.24 | 19.52 | 14.01 | 29.67 | 24.16 | 46.05 | 40.54 | 52.11 | 46.60 | 58.35 | 52.84 | 64.02 | 58.51 | 65.64 | 60.13 |
| Suraxani | Azerbaijan | 40.42 | 50.00 | 8.06 | 8.63 | 0.57 | 9.65 | 1.59 | 10.98 | 2.92 | 13.09 | 5.03 | 17.33 | 9.27 | 19.13 | 11.07 | 20.55 | 12.49 | 20.83 | 12.77 |
| Mastaga | Azerbaijan | 40.54 | 50.00 | 5.33 | 5.84 | 0.52 | 6.30 | 0.98 | 6.30 | 0.98 | 6.76 | 1.44 | 7.23 | 1.90 | 7.69 | 2.36 | 8.58 | 3.25 | 8.69 | 3.37 |
| Sabuncu | Azerbaijan | 40.52 | 49.97 | 7.30 | 7.44 | 0.14 | 7.69 | 0.40 | 8.14 | 0.84 | 8.72 | 1.42 | 9.18 | 1.88 | 9.62 | 2.32 | 10.51 | 3.21 | 10.95 | 3.65 |
| Galugah | Iran | 36.73 | 53.81 | 7.35 | 8.38 | 1.03 | 20.72 | 13.37 | 21.49 | 14.14 | 25.67 | 18.32 | 29.42 | 22.07 | 33.86 | 26.51 | 37.59 | 30.24 | 38.24 | 30.89 |
| Bandar-e Gaz | Iran | 36.77 | 53.95 | 1.97 | 3.08 | 1.10 | 18.64 | 16.67 | 21.30 | 19.32 | 25.51 | 23.53 | 32.12 | 30.15 | 37.81 | 35.84 | 42.84 | 40.87 | 43.46 | 41.48 |
| Kord Kuy | Iran | 36.79 | 54.11 | 7.31 | 9.40 | 2.09 | 23.14 | 15.83 | 27.21 | 19.90 | 32.25 | 24.94 | 39.35 | 32.04 | 47.93 | 40.62 | 52.86 | 45.55 | 54.57 | 47.26 |
| Sulak | Russia | 42.43 | 47.90 | 3.83 | 6.03 | 2.20 | 6.25 | 2.41 | 6.75 | 2.92 | 8.12 | 4.29 | 9.04 | 5.21 | 11.16 | 7.33 | 11.71 | 7.88 | 11.71 | 7.88 |
| Lagan | Russia - North basin | 45.39 | 47.35 | 18.30 | 20.39 | 2.09 | 59.91 | 41.60 | 71.83 | 53.53 | 72.12 | 53.82 | 72.53 | 54.23 | 72.95 | 54.65 | 130.43 | 112.13 | 140.80 | 122.50 |
| Zhanbay | Kazakhstan - North basin | 47.04 | 50.81 | 10.94 | 17.45 | 6.51 | 44.12 | 33.19 | 91.55 | 80.61 | 240.91 | 229.97 | 262.26 | 251.33 | 265.74 | 254.81 | 266.25 | 255.31 | 266.25 | 255.31 |
| Burynshik | Kazakhstan - North basin | 45.40 | 51.77 | 3.14 | 6.58 | 3.44 | 30.92 | 27.78 | 40.25 | 37.11 | 118.27 | 115.14 | 122.93 | 119.79 | 136.87 | 133.74 | 138.96 | 135.82 | 138.96 | 135.82 |
| Hazar | Turkmenistan | 39.42 | 53.10 | 0.00 | 0.36 | 0.36 | 0.72 | 0.72 | 1.14 | 1.14 | 2.31 | 2.31 | 4.03 | 4.03 | 7.73 | 7.73 | 10.40 | 10.40 | 11.96 | 11.96 |
| Esenguly | Turkmenistan | 37.46 | 53.97 | 7.59 | 8.77 | 1.19 | 15.26 | 7.67 | 21.63 | 14.04 | 30.97 | 23.38 | 38.14 | 30.55 | 43.79 | 36.20 | 48.54 | 40.95 | 49.41 | 41.82 |
| Sumqayit | Azerbaijan | 40.61 | 49.66 | 0.33 | 1.23 | 0.91 | 1.51 | 1.19 | 1.81 | 1.48 | 2.13 | 1.81 | 2.64 | 2.31 | 3.54 | 3.21 | 5.99 | 5.67 | 6.56 | 6.23 |
| Lankaran | Azerbaijan | 38.76 | 48.85 | 0.68 | 1.11 | 0.43 | 1.96 | 1.28 | 3.68 | 3.00 | 5.53 | 4.85 | 8.98 | 8.30 | 11.78 | 11.11 | 16.81 | 16.13 | 16.92 | 16.24 |

(continued)

| | | | | | Decline Scenario | | | | | | | | | | | | | | | |
|---|---|---|---|---|---|---|---|---|---|---|---|---|---|---|---|---|---|---|---|---|
| | | | | Sea level at -27.5m datum | 2.5m | | 5m | | 7.5m | | 10m | | 12.5m | | 15m | | 17.5m | | 18m | |
| Human Settlement | Country | Latitude | Longitude | Distance to shore, km | Distance to shore, km | Change in distance to shore, km | Distance to shore, km | Change in distance to shore, km | Distance to shore, km | Change in distance to shore, km | Distance to shore, km | Change in distance to shore, km | Distance to shore, km | Change in distance to shore, km | Distance to shore, km | Change in distance to shore, km | Distance to shore, km | Change in distance to shore, km | Distance to shore, km | Change in distance to shore, km |
| Atyrau | Kazakhstan - North basin | 47.09 | 51.91 | 27.10 | 33.90 | 6.80 | 60.76 | 33.66 | 83.21 | 56.11 | 286.78 | 259.68 | 290.46 | 263.36 | 295.37 | 268.27 | 295.93 | 268.83 | 295.93 | 268.83 |
| Sari | Iran | 36.57 | 53.06 | 23.97 | 23.97 | 0.00 | 24.28 | 0.31 | 24.71 | 0.73 | 25.14 | 1.16 | 25.76 | 1.79 | 26.56 | 2.59 | 27.20 | 3.23 | 27.55 | 3.58 |
| Babol | Iran | 36.56 | 52.67 | 17.38 | 17.38 | 0.00 | 17.63 | 0.25 | 17.93 | 0.55 | 18.32 | 0.94 | 18.72 | 1.34 | 19.23 | 1.85 | 19.75 | 2.37 | 19.97 | 2.59 |
| Amol | Iran | 36.47 | 52.36 | 20.06 | 20.06 | 0.00 | 20.13 | 0.07 | 20.41 | 0.35 | 20.67 | 0.61 | 20.99 | 0.93 | 21.31 | 1.25 | 21.56 | 1.50 | 21.66 | 1.60 |
| Gorgan | Iran | 36.84 | 54.44 | 35.54 | 36.28 | 0.75 | 44.79 | 9.26 | 48.75 | 13.22 | 54.47 | 18.93 | 61.66 | 26.12 | 71.70 | 36.16 | 77.39 | 41.86 | 79.44 | 43.90 |
| Makhachkala | Russia | 42.99 | 47.47 | 1.90 | 1.90 | 0.00 | 1.90 | 0.00 | 1.96 | 0.06 | 2.36 | 0.46 | 4.34 | 2.45 | 9.78 | 7.88 | 14.52 | 12.62 | 15.21 | 13.32 |
| Astrakhan | Russia - North basin | 46.35 | 48.08 | 55.00 | 65.97 | 10.98 | 119.28 | 64.28 | 149.68 | 94.69 | 157.18 | 102.18 | 168.53 | 113.53 | 174.49 | 119.49 | 218.14 | 163.15 | 218.71 | 163.71 |
| Rasht | Iran | 37.27 | 49.59 | 21.39 | 21.44 | 0.05 | 21.67 | 0.29 | 21.91 | 0.52 | 22.36 | 0.97 | 22.54 | 1.15 | 22.92 | 1.53 | 23.29 | 1.90 | 23.29 | 1.90 |
| Baku | Azerbaijan | 40.38 | 49.87 | 0.77 | 0.86 | 0.09 | 1.37 | 0.60 | 3.44 | 2.67 | 7.02 | 6.25 | 11.07 | 10.29 | 15.21 | 14.44 | 26.13 | 25.35 | 26.59 | 25.81 |
| Aktau | Kazakhstan | 43.65 | 51.17 | 2.35 | 2.35 | 0.00 | 2.35 | 0.00 | 2.35 | 0.00 | 2.42 | 0.07 | 2.78 | 0.43 | 3.07 | 0.72 | 3.75 | 1.40 | 3.88 | 1.53 |
| Bandar-e Anzali | Iran | 37.47 | 49.48 | 0.50 | 0.55 | 0.05 | 0.55 | 0.05 | 1.01 | 0.51 | 1.01 | 0.52 | 1.47 | 0.97 | 1.54 | 1.04 | 1.93 | 1.44 | 1.93 | 1.44 |
| Derbent | Russia | 42.06 | 48.29 | 0.00 | 1.62 | 1.62 | 1.95 | 1.95 | 2.09 | 2.09 | 2.41 | 2.41 | 3.40 | 3.40 | 4.19 | 4.19 | 4.68 | 4.68 | 4.85 | 4.85 |
| Kaspiysk | Russia | 42.89 | 47.62 | 4.86 | 4.86 | 0.00 | 4.86 | 0.00 | 4.86 | 0.00 | 4.86 | 0.00 | 4.86 | 0.00 | 5.62 | 0.76 | 7.65 | 2.79 | 7.95 | 3.09 |
| Chalus | Iran | 36.66 | 51.42 | 3.47 | 3.47 | 0.00 | 3.47 | 0.00 | 3.60 | 0.13 | 3.82 | 0.35 | 3.91 | 0.44 | 4.18 | 0.71 | 4.35 | 0.88 | 4.35 | 0.88 |
| Turkmenbashi | Turkmenistan | 40.03 | 52.96 | 2.42 | 5.18 | 2.76 | 5.89 | 3.47 | 14.26 | 11.83 | 15.03 | 12.61 | 15.60 | 13.17 | 16.47 | 14.04 | 17.29 | 14.87 | 17.35 | 14.92 |
| Izberbash | Russia | 42.56 | 47.87 | 0.00 | 2.21 | 2.21 | 2.44 | 2.44 | 2.73 | 2.73 | 2.99 | 2.99 | 3.27 | 3.27 | 3.42 | 3.42 | 3.55 | 3.55 | 3.64 | 3.64 |
| Behshahr | Iran | 36.69 | 53.54 | 13.33 | 20.85 | 7.53 | 21.69 | 8.36 | 22.30 | 8.97 | 23.40 | 10.07 | 25.51 | 12.19 | 27.76 | 14.43 | 30.19 | 16.86 | 30.75 | 17.42 |
| Langarud | Iran | 37.21 | 50.08 | 15.97 | 16.96 | 0.99 | 17.66 | 1.70 | 18.30 | 2.34 | 19.09 | 3.12 | 20.04 | 4.08 | 20.90 | 4.94 | 21.61 | 5.65 | 21.77 | 5.80 |
| Qaracuxur | Azerbaijan | 40.40 | 49.98 | 5.63 | 5.85 | 0.22 | 6.24 | 0.61 | 7.81 | 2.17 | 11.29 | 5.66 | 14.37 | 8.74 | 18.44 | 12.81 | 23.79 | 18.15 | 23.92 | 18.28 |

Supplementary Table 10. The change in distance to shore for human infrastructure by, or within, the Caspian Sea under different sea level decline scenarios.

| | | | | | Decline Scenario | | | | | | | | | | | | | | | | |
|---|---|---|---|---|---|---|---|---|---|---|---|---|---|---|---|---|---|---|---|---|---|
| | | | | | Sea level at -27.5m datum | 2.5m | | 5m | | 7.5m | | 10m | | 12.5m | | 15m | | 17.5m | | 18m | |
| Operator/ Location | Infrastructure type | Country | Latitude | Longitude | Distance to shore, km | Distance to shore, km | Change in distance to shore, km | Distance to shore, km | Change in distance to shore, km | Distance to shore, km | Change in distance to shore, km | Distance to shore, km | Change in distance to shore, km | Distance to shore, km | Change in distance to shore, km | Distance to shore, km | Change in distance to shore, km | Distance to shore, km | Change in distance to shore, km | Distance to shore, km | Change in distance to shore, km |
| BP | Gas Processing Plant | Azerbaijan | 40.15 | 49.47 | 0.00 | 0.45 | 0.45 | 0.81 | 0.81 | 1.52 | 1.52 | 3.58 | 3.58 | 5.07 | 5.07 | 16.87 | 16.87 | 24.09 | 24.09 | 27.16 | 27.16 |
| SOCAR | Gas Processing Plant | Azerbaijan | 40.27 | 49.69 | 0.00 | 0.00 | 0.00 | 1.54 | 1.54 | 3.14 | 3.14 | 4.13 | 4.13 | 6.21 | 6.21 | 10.69 | 10.69 | 27.51 | 27.51 | 30.11 | 30.11 |
| SOCAR | Oil Refinery | Azerbaijan | 40.37 | 50.01 | 2.91 | 3.73 | 0.82 | 5.48 | 2.57 | 7.01 | 4.10 | 9.73 | 6.82 | 12.29 | 9.38 | 16.09 | 13.18 | 25.49 | 22.58 | 25.81 | 22.90 |
| SOCAR | Oil Refinery | Azerbaijan | 40.32 | 49.85 | 0.00 | 0.00 | 0.00 | 0.65 | 0.65 | 0.77 | 0.77 | 2.69 | 2.69 | 5.09 | 5.09 | 8.95 | 8.95 | 30.86 | 30.86 | 32.23 | 32.23 |
| Dagestan Oil | Oil Refinery | Russia | 43.01 | 47.46 | 0.62 | 0.74 | 0.12 | 0.98 | 0.36 | 1.25 | 0.63 | 1.73 | 1.11 | 4.79 | 4.17 | 10.99 | 10.37 | 15.53 | 14.91 | 16.29 | 15.67 |
| KMG | Oil Refinery | Kazakhstan | 43.61 | 51.30 | 5.61 | 5.61 | 0.00 | 5.91 | 0.30 | 6.18 | 0.57 | 6.51 | 0.90 | 7.01 | 1.40 | 7.57 | 1.96 | 9.01 | 3.40 | 11.31 | 5.70 |
| Turkmenneftgas | Oil Refinery | Turkmenistan | 40.01 | 52.98 | 1.46 | 2.66 | 1.20 | 3.01 | 1.55 | 13.44 | 11.97 | 14.51 | 13.05 | 15.58 | 14.11 | 16.39 | 14.93 | 17.22 | 15.75 | 17.24 | 15.77 |
| National Iranian Oil Engineering & Construction Co | Oil Refinery | Iran | 36.81 | 53.49 | 6.69 | 6.69 | 0.00 | 7.66 | 0.98 | 8.64 | 1.96 | 9.57 | 2.88 | 11.32 | 4.63 | 13.64 | 6.95 | 16.05 | 9.36 | 16.62 | 9.94 |
| Lagan Port | Port | Russia | 45.38 | 47.45 | 9.91 | 13.78 | 3.86 | 54.76 | 44.85 | 67.17 | 57.26 | 67.40 | 57.48 | 67.83 | 57.92 | 68.27 | 58.35 | 125.24 | 115.32 | 135.96 | 126.04 |
| Makhachkala Sea Trade Port | Port | Russia | 43.00 | 47.48 | 0.65 | 0.65 | 0.00 | 0.65 | 0.00 | 0.87 | 0.22 | 1.16 | 0.52 | 3.27 | 2.63 | 9.05 | 8.41 | 13.73 | 13.08 | 14.44 | 13.80 |
| Dubandi Port | Port | Azerbaijan | 40.43 | 50.28 | 0.19 | 1.33 | 1.14 | 1.63 | 1.44 | 2.27 | 2.08 | 6.39 | 6.20 | 8.56 | 8.37 | 10.86 | 10.67 | 12.36 | 12.17 | 12.67 | 12.48 |
| Port Bandar-e Anzali | Port | Iran | 37.45 | 49.44 | 3.17 | 3.17 | 0.00 | 3.57 | 0.39 | 3.92 | 0.75 | 4.11 | 0.94 | 4.38 | 1.21 | 4.56 | 1.38 | 4.85 | 1.67 | 4.85 | 1.67 |
| Nowshahr Port | Port | Iran | 36.60 | 51.49 | 5.93 | 5.93 | 0.00 | 6.16 | 0.23 | 6.33 | 0.40 | 6.59 | 0.66 | 6.92 | 0.99 | 7.74 | 1.81 | 8.44 | 2.51 | 8.58 | 2.65 |
| Neka Port | Port | Iran | 36.82 | 53.27 | 2.94 | 2.94 | 0.00 | 3.49 | 0.55 | 4.05 | 1.10 | 5.13 | 2.19 | 6.73 | 3.79 | 8.56 | 5.61 | 9.82 | 6.87 | 10.11 | 7.17 |
| Okarem | Port | Turkmenistan | 38.01 | 53.84 | 1.53 | 2.12 | 0.60 | 3.94 | 2.42 | 10.89 | 9.37 | 22.27 | 20.75 | 37.32 | 35.80 | 57.04 | 55.52 | 64.80 | 63.28 | 67.14 | 65.62 |
| Hazar Port | Port | Turkmenistan | 39.34 | 53.22 | 0.00 | 0.45 | 0.45 | 2.22 | 2.22 | 6.21 | 6.21 | 9.69 | 9.69 | 16.13 | 16.13 | 19.37 | 19.37 | 22.33 | 22.33 | 22.69 | 22.69 |
| Kiyanly | Port | Turkmenistan | 40.16 | 52.78 | 0.68 | 1.53 | 0.85 | 2.24 | 1.56 | 3.01 | 2.32 | 4.29 | 3.60 | 5.10 | 4.41 | 6.05 | 5.37 | 6.87 | 6.18 | 7.03 | 6.35 |
| Kuryk Port | Port | Kazakhstan | 43.18 | 51.67 | 1.22 | 1.22 | 0.00 | 2.47 | 1.25 | 3.47 | 2.25 | 5.46 | 4.24 | 7.61 | 6.39 | 8.25 | 7.03 | 8.82 | 7.59 | 8.98 | 7.76 |
| Aktau Port | Port | Kazakhstan | 43.60 | 51.22 | 0.57 | 0.57 | 0.00 | 0.57 | 0.00 | 0.94 | 0.37 | 1.59 | 1.03 | 2.27 | 1.70 | 3.23 | 2.67 | 5.07 | 4.50 | 5.40 | 4.83 |
| Fort Shevchenko Port | Port | Kazakhstan | 44.54 | 50.26 | 0.00 | 0.00 | 0.00 | 0.72 | 0.72 | 1.19 | 1.19 | 3.31 | 3.31 | 5.56 | 5.56 | 7.25 | 7.25 | 12.64 | 12.64 | 12.98 | 12.98 |
| Turkmenbashi International Seaport | Port | Turkmenistan | 40.01 | 52.99 | 1.31 | 2.33 | 1.02 | 2.70 | 1.39 | 13.55 | 12.24 | 14.63 | 13.32 | 15.65 | 14.34 | 16.57 | 15.26 | 17.37 | 16.06 | 17.41 | 16.11 |
| Prorva Port | Port | Kazakhstan | 45.84 | 53.01 | 7.83 | 33.39 | 25.57 | 68.84 | 61.01 | 94.89 | 87.07 | 224.56 | 216.73 | 230.03 | 222.21 | 245.44 | 237.62 | 247.67 | 239.84 | 247.67 | 239.84 |
